# Carbohydrate degradation machineries in lichen fungal symbionts reveal distinct symbiotic footprints across *Ascomycota*

**DOI:** 10.64898/2026.07.28.741190

**Authors:** David Díaz-Escandón, Phillip Resl, Toby Spribille

## Abstract

Lichens—the archetypal symbiosis—have long been known for their nutritional relationship, in which the photoautotrophic partners subsidize the carbon needs of their fungal symbiont. Yet, this subsidiary framing obscures the fact that lichenization has evolved multiple times across different fungal lineages and involves a wide array of phylogenetically distinct photosynthetic symbionts and subsidy types. Here, we compiled and functionally annotated 309 fungal genomes—including 24 newly generated metagenomically assembled genomes—with 191 representing lichen fungal symbionts spanning all taxonomic classes with lichen symbioses in *Ascomycota*. We found that lichen fungal genomes consistently had fewer annotations than other fungi, except for CAZymes. Moreover, the enzymatic machinery of lichen fungal symbionts exhibits a distinct bimodal pattern, with some genomes maintaining large enzymatic repertoires, while others hold some of the smallest sets in ascomycotan fungi. This pattern closely aligns with their photobiont subsidiary molecules; lichens subsidized by their photobiont with the polyol erythritol possess large enzymatic repertoires compared to those that receive glucose, sorbitol, or ribitol. These retained enzymes are primarily related to carbon-harvesting functions, often streamlined as redundant functionalities in symbioses with a supplied carbon source. Our results suggest that lichens may have more than one fate for their carbon subsidies, rather than solely operating as nutritional symbioses.

## Background

Lichens are the outcomes of long-standing symbiotic relationships between a fungus and a photoautotrophic symbiont—the photobiont, be it cyanobacteria, green algae or, in some cases, brown algal stramenopiles^1^. Traditionally, this symbiosis has been considered to be a nutritional exchange, where the photobionts provide the fungi with a carbon source, while the fungus forms an interwoven hyphal body which serves as a medium for water capture and environmental protection^2,3^. For over 150 years, this carbon subsidy has been considered the core transaction that defines this symbiosis^4,5^. However, back in the 1960s, a series of experiments conducted by Smith and collaborators^6,7,8,9,10,11,12^ revealed that different photobionts release different photosynthates to the symbiosis—glucose in the case of cyanobacteria and a series of different acyclic polyols in green algae. During these experiments, a recurring result was that not all photosynthates share the same fate: some were incorporated into the central metabolism of fungal cells, while others easily washed away—not metabolically integrated—suggesting that the released photosynthates may play different roles in the symbiosis^13,14^. More recently, advances in genomics have shown that some fungi in lichen symbioses retain a large enzymatic machinery for degrading plant cell wall carbohydrates, suggesting the accessibility of external carbon sources from the symbiosis^15^. Despite these findings, the textbook definition has largely remained unchanged, although new hypotheses now account for the possibility of additional symbiont roles in osmoprotection and oscillating rewetting periods^1,2,3,16,17,18,19,20^.

Lichen symbiosis, as a lifestyle, has emerged multiple times throughout fungal evolution, occurring in both the fungal phyla *Ascomycota* and *Basidiomycota*^21^. Within *Ascomycota*, lichenization occurs in many independent lineages, with the highest diversity clustered in the class *Lecanoromycetes*^22^. Nonetheless, lichen symbioses are also found in other ascomycotan classes predominantly associated with heterotrophic lifestyles—including saprotrophs, and at times pathogens and parasites—such as *Arthoniomycetes*, *Eurotiomycetes*, *Dothideomycetes*, and, more recently, *Leotiomycetes*^22,23,24^. This seemingly typical pattern of lichen fungi sharing an evolutionary history with non-symbiotic fungi—as well as the radiation events associated with lichenization across *Ascomycota*—has been a subject of debate, with models ranging from single-origin lichenization events derived into multiple subsequent losses^25^, to numerous independent events with later acquisition of the lifestyle^26^. In more recent studies, it has been shown that lichenization also occurs in less flexible classes, such as *Lichinomycetes sensu lato*, a class where all taxa appear to be associated with a symbiotic lifestyle—including mycorrhizae, endophytes and yeast-like symbionts in beetle guts^27^.

In contrast to the fungal symbionts, the photosynthetic partners involved in lichen symbioses derive from completely independent origins, not restricted to symbiotic lifestyles^28,29,30^. They range from cyanobacteria to protists in *Stramenopiles* and chlorophycean green algae, belonging to the classes *Trebouxiophyceae, Chlorophyceae*, and *Ulvophyceae*. Among ascomycotan lichens, no photobiont group is shared across all fungal classes^1,28,29^. These nearly combinatorial interactions also involve a variety of different photosynthates entering the symbioses, including glucose, sucrose and trehalose released by cyanobacteria^14,31^; meanwhile, green algae release different polyols with varying carbon numbers: the four-carbon erythritol from trentepohlian-photobionts in *Ulvophyceae*, the five-carbon ribitol from *Trebouxiales*, and the six-carbon sorbitol from *Prasiolales*^1,2,13,32,33,34,35^. Notably, most of these molecules are also produced outside the symbiosis, with glucose and trehalose serving as primary carbon sources, while polyols primarily act as osmolytes during desiccation or osmotic stress^35,36,37,38^. Although the role of these photosynthates in symbiosis has largely been attributed to nutrition, experiments from the 1960s demonstrated that when incorporated into the symbiosis, these photosynthates exhibit different assimilation rates by their fungal symbiont, with some—such as erythritol, and to some degree ribitol—being largely non-incorporated into insoluble cellular components^13,39,40,41^. Today, while some of the anabolic pathways for the utilization of some of these algal derived polyols remain unknown, polyols—such as the six-carbon mannitol—are recognized to be the main carbon storage unit in fungi^42^.

This variability is not unique to lichen symbioses. Mycorrhizae—a nutritional partnership in which plant roots supply carbon-rich compounds and vitamins in exchange for fungal nitrogen and phosphates — evolved independently many times during fungal evolution^43^. Although the plant–fungus nature is universal, mycorrhizae are classified into different groups by their function and structure^44^. Comparative genomics have shown that arbuscular– and ectomycorrhizal genomes have reduced repertoires of carbohydrate active enzyme (CAZyme) machineries—especially plant cell wall degrading enzymes (PCWDEs)—while expanding their number of sugar transporters and invertases^45,46,47,48^.

In comparison, orchid– and ericoid-mycorrhizae maintained enriched PCWDE arsenals, resembling those from wood decayers and endophytes^48,49,50,51,52^. Comparatively, lichen symbioses mostly have been categorized based on their photobionts or morphological growth, not their symbiotic function. With the development of new molecular techniques, it has been shown that while some lichen fungi exhibit reductions in PCWDEs and sugar transporters, others maintain high numbers of PCWDEs and peroxidases^15,27,53^. However, these have been early approaches focusing on lichens from specific ascomycotan classes.

The predicted functional diversity of mycorrhizal fungi involved in different relationships with plants led us to hypothesize that lichen fungal genomes are shaped, over evolutionary timescales, by the identity of the sugars and sugar alcohols they take up from their photobionts. We hypothesized that lichen fungal genomes would exhibit distinct genomic profiles corresponding with the carbon subsidy and photobiont, shaped by the metabolic availability of carbon derived from the symbiosis and other non-nutritional exchanges. To test this, we first disentangled the effect of the different origins of the symbiosis from their various photoautotroph symbionts. We compiled all available genomes from lichen fungal symbionts across all ascomycotan taxonomic classes, along with non-lichenized fungal peers, and compared them as discrete units, distinct from their photobionts. Second, we compared the genomes of lichen fungal symbionts that receive different carbon subsidies. We integrated bibliographic data on photobionts and carbon subsidies with annotated genomic features, using genomic signatures as a proxy to validate the literature. Third, we compared lichen fungal CAZyme machinery profiles to identify common trait losses and compensatory adaptations associated with carbon scavenging. Our compilation of 309 ascomycotan genomes reveals multiple genomic signatures in lichen fungal genomes. Some profiles support the expected losses in CAZymes characteristic of a nutritional symbiosis. In contrast, others exhibit higher retention levels, suggesting that carbon continues to be acquired from sources other than the photobiont secretome and that carbon subsidies in lichens may serve functions beyond mere nutrition.

## Results

### Available data and new genomes

We analyzed a total of 309 fungal genomes, encompassing all taxonomic classes within the filamentous fungi in the ascomycotan subphyllum *Pezizomycotina* (308), along with one *Saccharomycetes* yeast as an outgroup (Table S1). Among these, 191 were lichen fungal genomes, including 130 assembled genomes and metagenomic-assembled genomes (MAGs) retrieved from the NCBI Assembly database or previously published (*Graphis scripta* in Resl et al. 2022), 38 lichen-derived libraries from the NCBI SRA database, and 23 newly produced MAGs in this research (Table S2). The lichen fungal genome set includes representatives from all the ascomycotan classes known to form lichen symbioses: *Arthoniomycetes* (12), *Dothideomycetes* (11), *Eurotiomycetes* (11), *Lecanoromycetes* (133), *Leotiomycetes* (1), *Lichinomycetes* (19), as well as species classified as *incertae sedis* (4).

From the 191 lichen fungal genomes—representing 183 species—we derived photobiont information from the literature, resolving at least to the genus level for 91 species. In the remaining 92 species, the literature describes the photobionts in broad terms such as ‘green algae’ or ‘chlorococcoid’, and in a few cases more specifically as ‘trentepolioid’ or ‘trebouxioid’ (Table S3). Of the 121 genera represented in our set, only six—*Caeruleum*, *Piccolia*, *Pycnora*, *Strangospora*, *Thelenella,* and *Felipes*—lack genus-level photobiont identification, with bibliographic sources referring to their photobionts simply as ‘chlorococcoid’, ‘trebouxioid’, ‘trebouxiophycean’, or in the case of the genus *Felipes*, ‘trentepohlioid’. We validated and updated the photobiont identifications in all the metagenomic-derived genomes with available ITS and rbcL sequences (Table S3).

Our lichen fungal genomes set included symbioses with 20 different photobiont genera, including 12 genera in *Chlorophyta* (green algae) and eight in *Cyanobacteria*. The chlorophytan symbionts belong to two classes and four orders: *Trentepholiales* in *Ulvophyceae* and *Trebouxiales*, *Prasiolales*, *Watanabeales* in *Trebouxiophyceae*, along with three *incertae sedis* genera*: Elliptochloris, Coccomyxa* (*Elliptochloris*-*Coccomyxa* clade), and *Leptosira*.

To contrast the different symbiotic relationships that exist between the lichen fungi and their different photobionts, we surveyed and compiled all literature-reported information on potential photosynthates produced or released by the photobionts either in the free-living or symbiotic state. We found experimental data referencing major secreted sugars or sugar alcohols for 11 out of the 12 green algae genera identified in this study. The only genus with no information was *Leptosira*, which is paraphyletic to the orders *Watanabeales* and *Trebouxiales*^54^, both commonly associated with ribitol secretion (Table S4). Among the most common polyols recognized in these symbioses, erythritol has been identified in *Trentepohliales*, ribitol in *Watanabeales* and *Trebouxiales*, and sorbitol in *Prasiolales*. For cyanobacterial symbionts we found experimental references covering three genera within the *Nostocales* and one genus in *Pleurocapsales*. Collectively, as *Cyanobacteria,* they are known to secrete glucose, followed by the non-reducing disaccharide trehalose (Table S4).

### Lichen fungal genomes are relatively smaller than other ascomycotan fungi

Lichen fungal genomes exhibit a heterogeneous range in size, spanning from some of the smallest genomes in filamentous *Ascomycota* (14.8 Mbp in the class *Lichinomycetes*) to others considerably exceeding the average genome size for the group (up to 131 Mbp in the order *Peltigerales, Lecanoromycetes*). On average, lichen fungal genomes have a genome size of 34 Mb (SD: 13.9 Mb, n = 191), a smaller genome size (Wilcoxon rank-sum test: *p* = 0.01) than the average non-lichen fungal genome of 38.5 Mb (SD: 20.35 Mb, n = 118). Regarding gene content, lichen fungal genomes contain an average of 9,008 genes (SD: 1,392), significantly fewer than the 10,259 genes (SD: 2,609) found in non-lichen fungal genomes (Wilcoxon rank-sum test: *p* < 0.001), with more than 1,000 fewer genes. The GC content in lichen fungal genomes was also slightly lower, with an average of 47.2% (SD: 4.83) compared to 48.7% (SD: 4.20) in non-lichen fungal genomes (Wilcoxon rank-sum test: p = 0.003). The most significant difference across lichen fungal genomes was in the number of tRNAs (Wilcoxon rank-sum test: *p* < 0.001), with an average of 56 copies (SD: 42). In contrast, the non-lichen fungal genomes averaged 177 copies (SD: 330). Despite the differences, the data density distribution is remarkably similar across genome size, gene number, and GC content, with most of the variation explained by heterogeneous sampling and sample sizes (Fig. 1A–H, Table S5).

**Figure 1.**
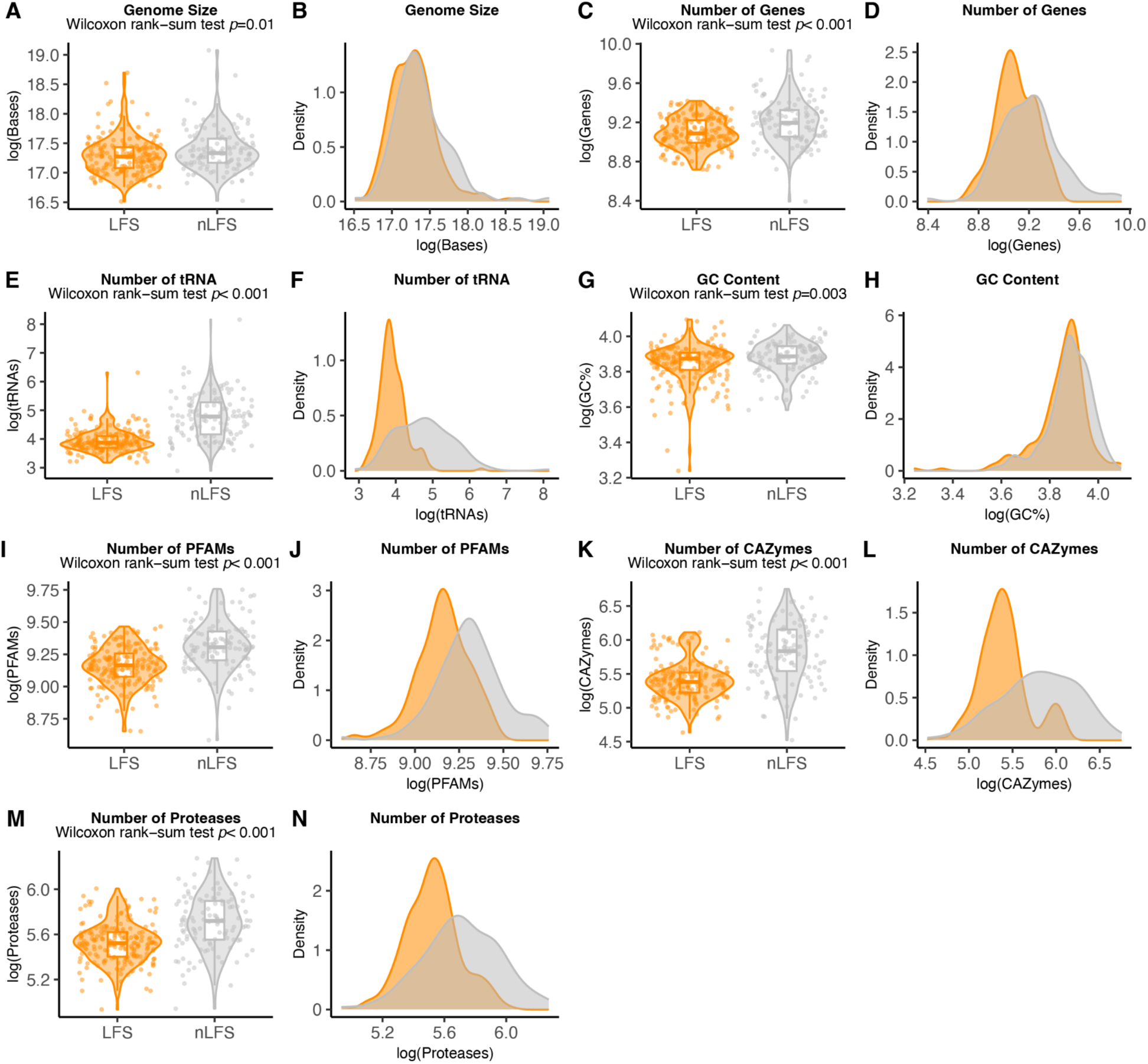
Genomic features comparisons and density plots of 309 fungal genomes. In orange 191 lichen fungal symbiont genomes (LFS) and in gray 118 non-lichen fungal symbiont genomes (nLFS). (A–B) Data based on genome size, with log-transformed number of base pairs per genome. (C–D) Data based on the log transformation of the number of predicted genes. (E– F) Data based log transformation of the number of copies of tRNAs per genome. (I–J) Data based on the log transform count of Pfam annotations. (K–L) Data based on the log transformation of CAZymes annotated proteins. (M–N) Data based on the log transformation of protease annotations. Boxplots integrated with violin distribution; each dot represents a genome. Density plots with log-transformed data. All comparisons included a Wilcoxon rank-sum test, and only *p-values* above 0.001 are displayed.

Next, we compared the gene annotations between lichen and non-lichen fungal genomes (Fig. 2I–N, Table S5). We found significant differences in the number of total protein family (PFAM) annotations, with lichen fungal genomes consistently having fewer, averaging 9,614 (SD: 1,569) total PFAMs within 3,557 (SD: 220) protein families. In contrast, the non-lichen fungal genomes averaged 11,285 (SD: 2147) total pfams, from an average of 3832 (SD: 192) protein families. Similarly, annotations for proteases and carbohydrate-active enzymes (CAZymes) of lichen fungal genomes hold significantly smaller sets than non-lichen fungal genomes, averaging 254 (SD: 45) annotated proteases and 231 (SD: 71) annotated CAZymes. In contrast, non-lichen fungal genomes averaged 312 (SD: 77) and 366 (SD: 154), respectively. Contrary to the average genomic features mentioned above, the data density distribution was highly heterogeneous, with the broadest distribution in CAZyme annotations. While PFAM and protease annotations retain a mostly unimodal pattern, CAZyme distribution exhibits two distinct peaks, spanning from the lowest numbers to above the non-lichen fungal genome average; however, it was not significantly different than a unimodal distribution (Hartigan’s Dip Test *p* = 0.9).

**Figure 2.**
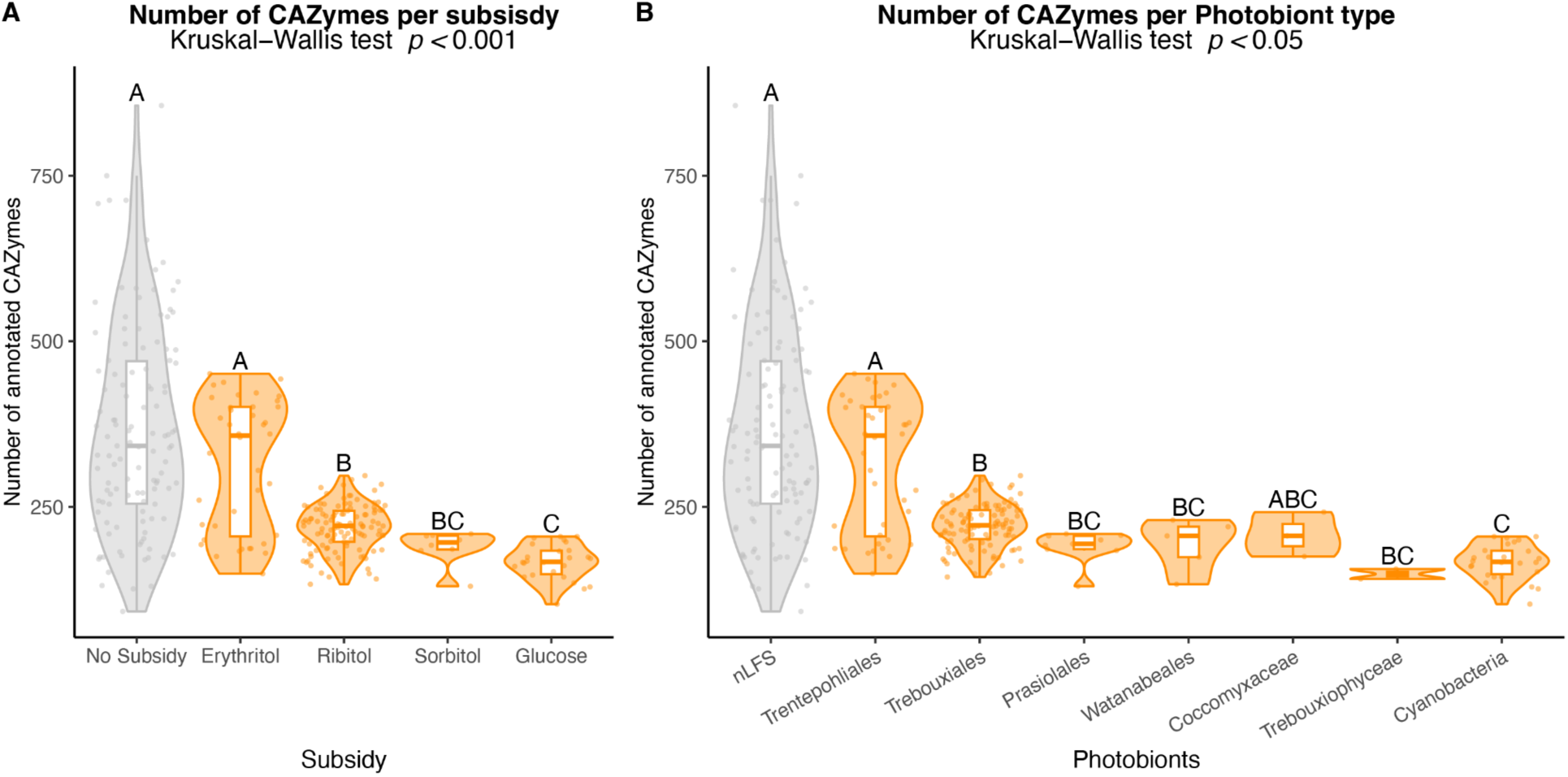
CAZyme annotation comparisons across known lichen subsidy molecules and photobiont clades in 307 fungal genomes. In orange 189 lichen fungal symbiont genomes (LFS), after excluding lichen with unknown symbionts, and in gray 118 non-lichen fungal symbiont genomes (nLFS). Groups based on the multiple comparison Dunn’s Test with *p* < 0.05. (A) Based on the number of annotated CAZymes and grouped by literature-reported lichen subsidy molecules. (B) Data based on number of annotated CAZymes and group by literature-reported photobionts in lichens, the clades *Coccomyxaceae* and *Trebouxyphyceae* account for three and two samples respectively

Additionally, we compared the genomes of each fungal taxonomic class containing lichen symbioses (with two or more representative species) between lichen fungal and non-lichen fungal genomes (Table S6–S7). We found significant differences sparsely distributed within classes, with lichen fungal genomes having fewer number of annotations in all cases. In the case of tRNAs, lichen fungal genomes in the *Eurotiomycetes* and *Dothideomycetes* hold significantly lower counts than their non-lichenized peers (Wilcoxon rank-sum test: *p* < 0.01). In contrast, non-lichen fungal genomes in *Lichinomycetes* and *Lecanoromycetes*—although with slightly higher counts—show no significant differences from the lichen peers. PFAMs were significantly different between lichen and non-lichen fungal genomes in *Eurotiomycetes* and *Lichinomycetes* (Wilcoxon rank-sum test: *p* < 0.05), with non-lichen fungi holding higher counts of PFAMs. In contrast, *Lecanoromycetes* and *Dothidiomycetes* exhibit no significant differences. In proteases, non-lichen fungal genomes hold significantly higher annotation than their lichen peers in *Eurotiomycetes* and *Lecanoromycetes* (Wilcoxon rank-sum test: p < 0.05), with no significant differences among *Dothidiomycetes* and *Lichinomycetes*. CAZymes hold the highest variability within taxonomic classes with significant higher counts in non-lichen fugal genomes compared to lichen fungal genome in all classes—*Eurotiomycetes*, *Lecanoromycetes* and *Lichinomycetes* (Wilcoxon rank-sum test: p < 0.05)—, except *Dothidiomycetes* with non-significant differences.

### Subsidy types define, at least two, distinct lichen enzymatic profiles

Next, we investigated whether the bimodal distribution in CAZyme repertoires is attributable to the different subsidies or the different photobionts. To do this, we compared the CAZyme machinery within lichen fungal genomes by grouping them according to their photobiont clades and the type of subsidy molecules released by their corresponding photobionts. We found that, in both scenarios (subsidies and photobionts), there were significant differences between the groups (Kruskal-Wallis, *p* < 0.001). More specifically, lichen fungi associated with cyanobacteria or glucose-subsidized photobionts hold the smallest CAZyme repertoires, averaging 167 gene annotations (SD: 27) (Fig. 2, Table S8–S9). In contrast, trentepohlian or erythritol-subsidized lichen fungal genomes tend to have larger CAZyme repertoires, but display high variability, with a standard deviation four times larger than that of other lichen fungal genomes. On average, these genomes had 315 gene annotations (SD: 99) and a median of 358 annotated genes, slightly exceeding that of non-lichen fungal genomes (median: 342) (Table S8–S9). In the trebouxiophycean orders, which include the ribitol– and sorbitol-subsidized lichen fungal genomes, a slight difference is observed. Genomes of lichen fungi that receive sorbitol, mainly from the algal order *Prasiolales*, have a slightly lower number of CAZymes (mean:189, SD:26), whereas ribitol-subsidized lichen fungal genomes represented by *Trebouxiales*, *Watanabeales* and *Coccomyxaceae* exhibit moderately larger CAZyme counts (mean:220, SD:33).

To identify shared profiles among lichen fungal genomes, we wanted to know how the genomes grouped across different subsidies and photobionts. To do this, we used Dunn’s Test for multiple comparisons separately on all lichen fungal symbiont genomes categorized by photobiont taxonomy and received subsidy, respectively. In each test, we identified three distinct statistical levels (Fig. 2A, Table S8). The first level included trentepohlian or erythritol-subsidized lichen fungal genomes, which grouped with non-lichen fungal genomes or non-subsidized genomes. The second level consisted of ribitol-subsidized genomes from the *Trebouxiales* clade. The third level includes genomes associated with cyanobacteria or glucose-based subsidies. An intermediate level, characterized by sorbitol-subsidized genomes, emerged between ribitol– and glucose-subsidized genomes; however, this distinction became less pronounced in the photobiont-based analysis, likely due to additional subdivision within the ribitol-subsidized genomes (Fig. 2B, Table S9).

Moreover, in the multiple comparisons, the two peaks in CAZyme distribution previously observed in lichen fugal genomes (Fig. 1L) persist, but only in the trentepohlian or erythritol-subsidized lichen fungal genomes (Fig. 2). We tested this group for multimodality and confirmed a significant bimodal distribution (Hartigan’s dip test, *p* = 0.002). This bimodality places one subset of the erythritol-subsidized lichen fungal genomes closer to the ribitol group, with a smaller number of annotated CAZymes. In contrast, the remaining trentepohlioid lichen genomes are closer to the average number of CAZymes for non-lichen fungal genomes.

### CAZyme composition varies among subsidies

To analyze the distinctive profiles produced by the different types of subsidy molecules, we examined the CAZyme annotation composition in terms of CAZyme classes. For this analysis, we partitioned the abundance of CAZyme annotations by CAZyme class in lichen-derived fungal genomes only and aggregated them in raw counts and relative ratios. Erythritol-subsidized genomes exhibit the highest variability, including the highest numbers of annotated CAZymes in all classes (Fig. 3A, Table S10). In terms of composition, glycoside hydrolases (GH), which break down glycosidic linkages, are the most abundant CAZymes across all groups. Their prevalence ranges from 49% (average 155) in erythritol-subsidized fungal genomes to 37.8% (average 63) in glucose-subsidized ones (Fig. 3B, Table S10). In contrast, polysaccharide lyases (PL), which cleave specific glycosidic linkages in acidic polysaccharides, are the least abundant, ranging from 0.9% in erythritol– and sorbitol-subsidized genomes to 0.5% in ribitol-subsidized lichen fungal genomes. Glycosyl transferases (GT), which form glycosidic bonds, were highly conserved across all groups, with raw counts ranging from an average of 67 annotations in erythritol and ribitol to the lowest averages of 58 and 57 in sorbitol and glucose, respectively. However, in relative ratios, GTs comprise 34.8% of the enzymatic machinery in glucose-subsidized genomes, compared to 19.5% in erythritol-subsidized lichen fungal genomes, a function of the higher number of non-GT CAZymes in the latter.

**Figure 3.**
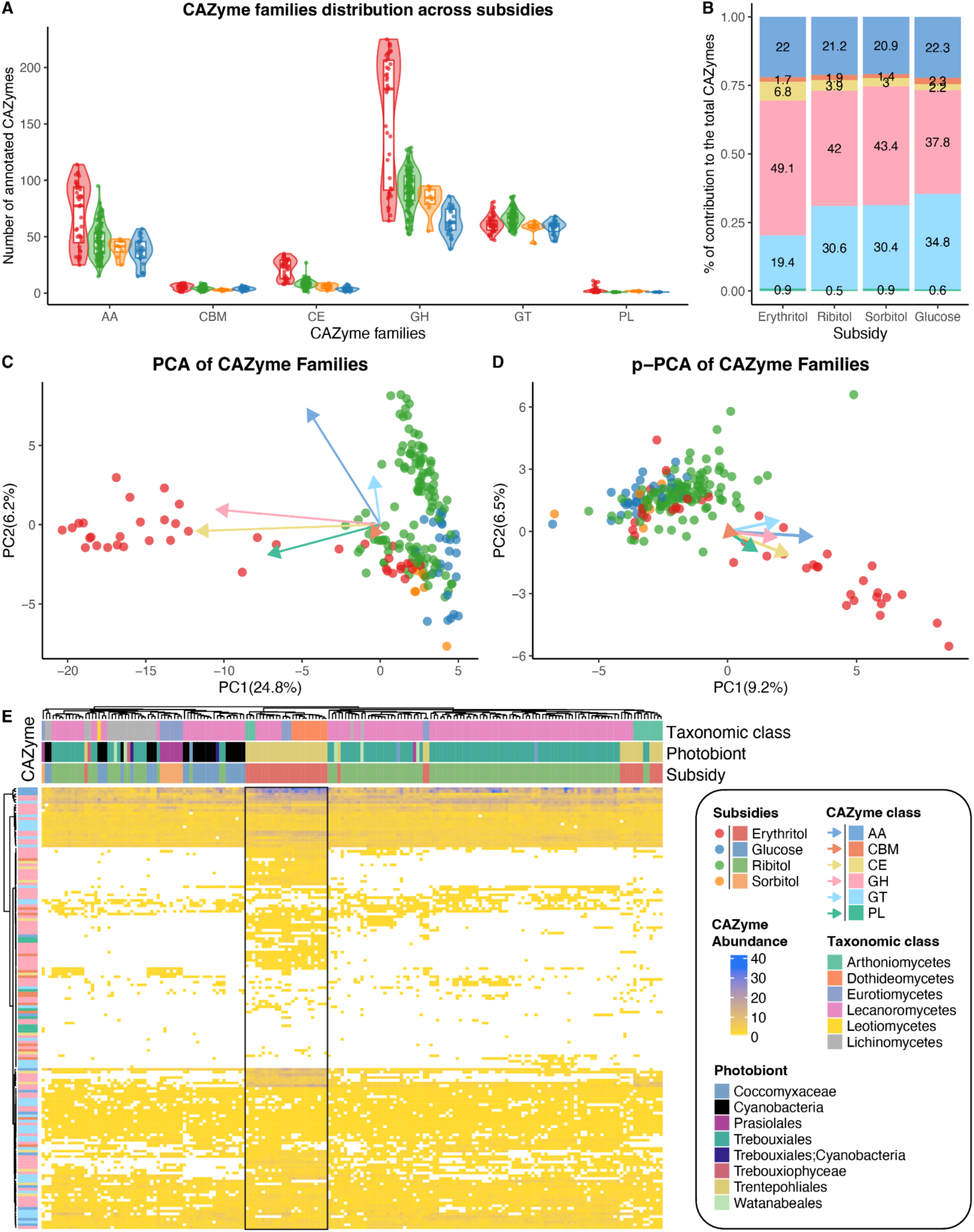
CAZyme family abundance and composition across different subsidy types in 189 lichen fungal genomes. (A) Barplot with density distribution based on annotations from 162 CAZyme families, split across CAZyme classes, colored by subsidy molecules. (B) Local proportion of CAZymes per subsidy based on annotations from 162 CAZyme families, where contribution is the mean of each CAZy class divided by the sum of all CAZy class means per subsidy. (C) PCA based on 162 CAZyme families, arrows show the median loading contribution per CAZyme class, scaled by the maximum dimensions of the principal components. (D) Phylogenetically corrected PCA (pPCA) based on 162 CAZyme families, arrows show the median loading contribution per CAZyme class scaled by the maximum dimensions of the principal components. (E) Heatmap of CAZyme annotations per lichen fungal genome along subsidy, photobionts and taxonomic class, based on 162 CAZyme classes, where each rows represent a CAZyme family and each column a lichen fungal genome, each cell represents CAZyme abundance, white for absence, and colour transition from annotation abundance from yellow (1) to blue (40).

To profile the enzymatic machinery associated with different subsidies, we analyzed the contribution of each CAZyme class and family across these subsidies. To achieve this, we performed a principal component analysis (PCA) using all CAZyme families from all lichen genomes and grouped them by subsidy (Fig. 3C). The CAZyme profiles formed two distinct clusters split across principal component 1 (PC1): with most of the erythritol-subsidized genomes on the left side, while glucose-, sorbitol-, and ribitol-subsidized genomes formed a cluster on the right side. Along PC2, the data aligned such that ribitol-subsidized genomes disperse across the entire axis, whereas glucose-, erythritol– and sorbitol-subsidized genomes tended to cluster towards the top.

Additionally, to evaluate the contribution of CAZymes classes to the clustering distribution, we analyzed the PCA loading weights for each CAZyme class (represented as arrows in Fig. 3C, Table S11–S12). To summarize the loadings, we calculated the median of the PCA loadings for each principal component across all CAZyme families within each CAZyme class (Fig. 3C). Carbohydrate Esterases (CE), which catalyze the release of acyl– or alkyl-groups linked by ester bonds to carbohydrates, and GHs are the main contributors of PC1, which correlates with the clustering of erythritol-subsidized lichen fungal genomes. Additionally, PLs family load contributed substantially to PC1, despite their overall low abundance in the total CAZyme repertoire—the lowest contribution to PC1 comes from the loads of carbohydrate-binding modules (CBM) and GTs. Variation along PC2 mainly relies on the redox enzyme class of auxiliary activities (AA), which partially contributed to PC1, associated with the split between the ribitol-subsidized lichen fungal genomes.

We next considered the possibility that some or all of the detected CAZyme richness patterns could be attributed to the phylogenetic relatedness of the sampled genomes or to evolutionary conservatism. To test this, we constructed a species tree based on 1387 gene trees from shared single-copy orthologs identified from AscomycotaDB10 (BUSCO), including only those with a mean bootstrap support of 80% or higher per gene to ensure stability (Fig. S1). Using this species tree, we test phylogenetic signals along PC1 and PC2. Pagel’s λ indicates a strong phylogenetic signal along PC1 and PC2 of 0.98 and 0.96, respectively (*p* < 0.0001), where λ=1 corresponds to the expectation of Brownian Motion in a continuous trait. To account for this effect, we conducted a phylogenetically corrected principal component analysis (p-PCA) in conjunction with the PCA to evaluate the contribution to the variance (Fig. 3D, Table S13–S14). The p-PCA Pagel’s λ reflected a considerable reduction at 0.63 and 0.46, respectively, for PC1 and PC2, while still carrying some phylogenetic signal. The primary division between the cluster formed by most of the erythritol-subsidized genomes and other subsidies remained evident but now appears along the diagonal across PC1 and PC2. In the p-PCA, the first two components account for 15.7% of the observed variance, compared to 31% in the standard PCA, suggesting that phylogenetic relatedness may contribute substantially to the observed variance. Nonetheless, the main clustering patterns and the load contributions from CAZyme families remained similar, indicating that the significant trends observed at the clustering were persistent.

Following, we created a heatmap of CAZyme family abundance against fungal taxonomic classes, photobionts, and subsidies to examine general guilds or profiles across the lichen fungal genomes (Fig. 3E, Table S15). We detected a block of genomes with similar CAZyme family profiles linked to erythritol and trentepohlian photobionts, characterized by a higher abundance of CAZyme families, as well as CAZyme families not shared by other groups (indicated with a box in Fig. 3E). This profile was independent from the taxonomic class, including *Arthoniomycetes*, *Dothideomycetes*, *Eurotiomycetes*, and *Lecanoromycetes*. However, it was not the only profile reflected in erythritol subsidies, with some occurrences indistinguishable from those of ribitol, sorbitol, or glucose.

### Case by case, CAZyme family contribution

Since the clustering and profile signals among different lichen fungal genome groups were disproportionately affected by specific CAZyme families, we analyzed each CAZyme family individually, grouping them by the different substrate molecules (Fig. 3E). We used the non-parametric Kruskal-Wallis (K-W) test, along with generalized linear models (GLMs), using on a case-by-case basis either Poisson or negative binomial distributions. Out of the 162 CAZyme families shared between lichen fungal genomes, between 122 and 105 were significantly different among the subsidies (K-W test: BH corrected *p* < 0.05; Likelihood Ratio Test: BH corrected *p* < 0.05). One hundred and four (104) families were significantly different across both statistical approaches. Of these, 96 occurred in at least 5% of the lichen fungal genomes, which matches the minimum expected representation of the smallest subsidy group and removes the noise of local maxima produced by specific families or genera.

Carbohydrate esterases (CE), exhibited the highest percentage of significant differences, with 10 out of 13 families exhibiting significant variation. This was followed by glycoside hydrolases (GHs), with 53 out of 77, CBMs, with 10 out of 15 and AAs with eight out of 13. Glycosyl transferases (GTs) and polysaccharide lyases (PLs) displayed the least variability, with 12 out of 37 and 3 out of 8 families showing significant differences between the subsidies.

We conducted a series of post hoc analyses, including Dunn’s test for the Kruskal-Wallis results and Estimated Marginal Means for the GLMs, to assess each CAZyme family grouping among different subsidies. In 36 out of 96 CAZyme families with significant differences, erythritol-subsidized genomes formed a distinct group, while those subsidized by ribitol, sorbitol, and glucose consistently clustered together at the same level (Fig. 4, Table S16–S19). The second most frequent pattern, accounting for 10 CAZyme families, introduced a third level of distinction: erythritol-subsidized genomes remained in a separate category, ribitol-subsidized genomes formed a second level, glucose-subsidized genomes constituted a third level, and sorbitol-subsidized genomes were indistinguishable from those at the ribitol and glucose levels. Additional patterns included cases where glucose and sorbitol formed a group, while other sugars remained at independent levels (nine families); erythritol and ribitol formed a group, while sorbitol and glucose formed another group (five families).

**Figure 4.**
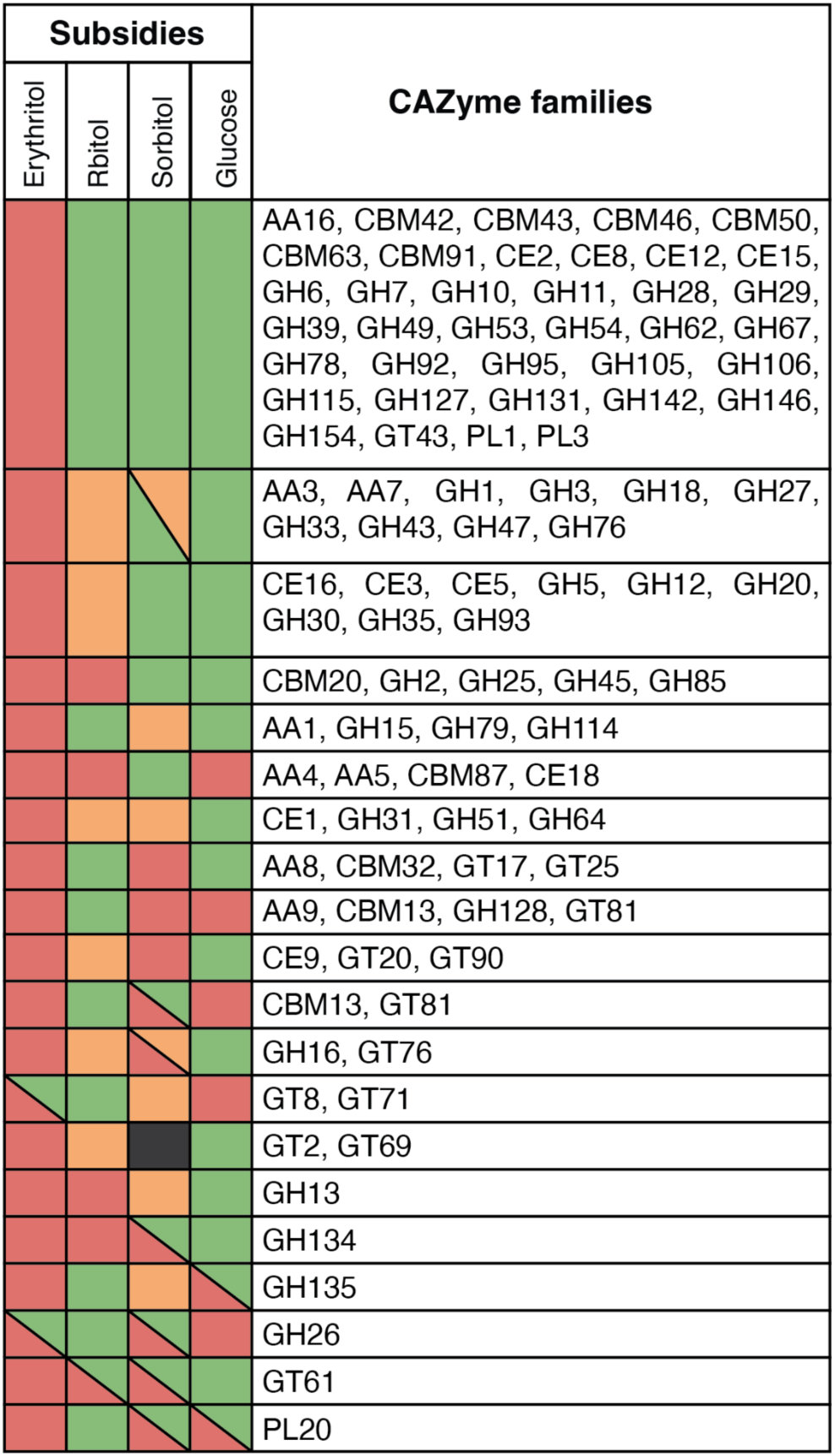
Grouping of 96 CAZyme families per subsidy from 189 lichen fungal genomes. Colours show Dunn’s post hoc grouping (*p* < 0.05); dark gray indicates unresolved or “ABC” classifications. Groups are not directly comparable but consist of level assignments irrespective of their directionality.

Eight CAZyme families were found exclusively in erythritol-subsidized lichen fungal genomes, with no occurrences in other groups. These included pectin and pectate lyases (PL1 and PL3), the carbohydrate-binding module CBM46, arabinofuranosidases (GH62), glucuronosidases (GH67), fuconosidases (GH29), the family GH39 of xylosidases, which was almost exclusive to the class *Dothideomycetes* within erythritol-subsidized genomes, and the carbohydrate esterase family CE15, which hydrolyzes lignin-carbohydrate ester linkages. Additionally, the families of glycosyltransferases GT81 and GT31 were exclusive to the ribitol-subsidized genomes. However, GT31 occurred in fewer than 10% of the genomes surveyed for the group, with no significant differences after adjustment for sample size (BH-corrected *p* > 0.05). We found no CAZyme families exclusive to sorbitol– nor glucose-subsidized genomes.

Furthermore, eight CAZyme families exhibited significant changes in abundance in erythritol-subsidized genomes with more than double the average number of annotations compared to those with other subsidies (Table S18–S19). The family GH28 of polygalacturonases averaged six annotations for erythritol-subsidized lichen fungal genomes, but it was nearly absent in all other groups. Similarly, the family GH92 of ɑ– mannosidase, which cleaves ɑ-mannose, displayed an average of four annotations for erythritol-subsidized genomes, one in ribitol-, and zero in sorbitol– and glucose-subsidized genomes—a ratio also observed for the family of arabinofuranosidases GH43. Additional CAZymes that more than doubled their annotation counts in erythritol-subsidized genomes included xylan esterases (CE5), acetylesterases (CE16), xylanases (GH11, nearly exclusive to erythritol-subsidized genomes), galactosidases (GH35), and β– glucuronidase (GH79).

Other CAZyme families highly representative of erythritol-subsidized genomes, but not exclusively found in them, include: multicopper oxidases (AA1), oxidoreductases (AA3) glucooligosaccharide oxidases (AA7), lytic cellulose monooxygenases (AA16), xylan-binding modules (CBM42 and CBM91), glucan– and cellulose-binding modules (CBM43 and CBM63), acetylesterases (CE2, CE3, and CE12), pectin methylesterases (CE8), and esterases interacting with diverse substrates in CE1 and CE5. Among the glycoside hydrolases they include cellulases (GH5, GH6, GH7 and GH12), fucosidases (GH95), glucanases (GH49, GH64, GH93, GH106 and GH131), glucuronidases (GH115), glucosidases (GH15), acetylglucosaminidases (GH20), glycosidases (GH27 and GH31), exosialidases (GH33), enzymes acting on hemicelluloses (GH51), xylanases (GH10), mannanases (GH76 and GH146), arabinofuranosidases (GH54, and GH127), mannosidases (GH47), polygalactosaminidases (GH114), pectin-degrading enzymes (GH53, GH78, and GH105), multifunctional families (GH30) and families with unknown functionality (GH142 and GH154).

These results, seemingly biased to erythritol-subsidized genomes, underscore how different these genomes are compared to the ones related to other subsidies. The only CAZyme families missing in erythritol compared to those found in genomes of fungi that receive other subsidies include the lytic polysaccharide monooxygenase family AA14, the galactosyltransferases family GT31, the mannosyltransferase family GT81 and the mannose– and xylan-binding module family CBM13. Nonetheless, none of these families are highly representative of any other subsidy. Additionally, the lysin-binding motif family CBM50—a known effector in plant pathogens—was detected in a single genome in the erythritol-subsidized group, whereas it was sparsely present but significantly more common in genomes receiving other subsidies.

Ribitol-subsidized genomes also form their own category. Specifically, the family of β-1,3– glucanases GH128 is the only case where a family significantly exceeds the number of annotations compared to the other subsidies. Additionally, the family abundance of the acetylesterases CE3 and CE16, together with the esterase family CE5, the endoglucanase family GH12, and the cellulase family GH5, form a distinct level below erythritol-subsidized genomes, but significantly different from other subsidies. In contrast, for the cellulose family AA9, ribitol-subsidized genomes exhibit the smallest set of annotations, even lower than those observed in sorbitol– and glucose-subsidized genomes.

In sorbitol-subsidized genomes, the vanillyl-alcohol oxidase AA4 was completely absent, while it was prevalent across all the other subsidies. The starch-binding family CMB20 was absent; however, the sparse occurrences across other subsidies yielded mixed results on the post hoc tests (Table S17–S19). Some groups exhibiting low numbers of annotations compared to the other subsidies include the multicopper oxidases AA1, with the lowest abundance of all, and AA5, as well as the glycosyltransferases GT8 and mannosyltransferase GT71. Sorbitol genomes also exhibited the highest number of annotations from the α-amylase family GH13 (averaging at 11), almost double that of all other subsidies. Other families with higher number of annotations include the family of pectin esterases CE18, the galactosaminogalactan-binding module CBM87, the family of inverting α-glycosidases GH15, and families of polygalactosaminidases and galactosaminidases GH114 and GH135. The family of β-glucuronidases, GH79, forms a distinct level below erythritol and above glucose and ribitol.

Glucose-subsidized genomes exhibited no CAZyme families present in higher abundance than in genomes of lichen fungi that received any other subsidy. Notably, the family of N-acetylglucosamine-6-phosphate deacetylases, CE9, which is involved in sugar metabolism and peptidoglycan recycling, was completely absent, whereas it occurs sparsely across all the other subsidy types. The family of β-glucosidases and β– galactosidases GH1 was also absent, as well as in sorbitol-subsidized genomes. Additionally, genomes of glucose-subsidized fungi contain the lowest number of annotations for some families highly represented in erythritol-subsidized genomes, including AA3, AA7, CE1, GH31, GH51, and GH64. In the family GT20 involved in trehalose-6-phosphate catalysis and the family of xylosyltransferases GT90, the glucose subsidy type exhibits the lowest number of annotations, below that of ribitol-subsidized genomes, with lower numbers also occurring in the families of α-glucoside linkages GH13 and the family of transglycosylases GH16.

## Discussion

Like many symbiotic relationships, lichen fungi are continuously exposed to selection pressures or redundancies within their symbiosis, leading to trait losses and compensatory adaptations^55^. The nature of their symbiosis—whether it is nutritional, osmo-protective, or with diverse photobionts—directs the trajectory of their genome evolution. Here, the different genome signatures observed suggest that ‘lichen symbiosis’ is an umbrella term encompassing multiple functional roles. But to what extent do these differences define unique symbiotic strategies? Or do they originate from a common symbiotic foundation, diverging through specialization with different photobionts or niche adaptations?

### Are the genomic signatures related to a single kind of symbiosis?

Lichens have been described as the archetypal nutritional symbiotic relationship, where the fungal symbiont benefits from photosynthates produced by a photoautotrophic partner. However, it remains debated whether this nutritional—yet undemonstrated— exchange represents the defining feature of the symbiosis or if additional functions are involved^1^. With the advent of comparative genomics, researchers have gained new tools to test and profile symbiotic relationships at the genomic level, revealing both short-term and long-term consequences to genome evolution in symbioses^15,27,48,53^.

Nutritional symbioses in the fungal kingdom are widely studied, and evidence indicates that substrate interactions—or carbon subsidies—play a crucial role in genome evolution^44,56,57^. Mycorrhizal symbioses across the fungal kingdom are carbon-subsidized and share a common genomic profile. This profile characteristically comes with a reduced enzymatic arsenal—mainly plant cell-wall-degrading enzymes (PCWDEs)—and an accumulation of transposable elements (TEs) and repetitive elements, resulting in larger genomes compared to their close non-symbiotic relatives^48,56,58^. Two main mechanisms could explain this common reduced signature: functional redundancy, which leads to relaxed selection where the fungi no longer rely on saprophytically harvested carbon, as carbon is subsidized^55,59^; and host-symbiont interactions, which select for the minimum enzymatic toolset required to remodel cell walls while minimizing host adverse responses^52^. That said, not all mycorrhizae share the same profile. Ericoid mycorrhizae and orchid mycorrhizae, both originating from multiple independent sources, possess extensive enzymatic arsenals and arguably distinct lifestyles^48,49,52^.

Recent genomic profiling of lichen fungal symbionts has revealed a similar reduction in enzymatic machinery—similar to carbon-subsidized mycorrhizal symbioses—but without the typical mycorrhizal genome expansion. The first attempt to characterize the CAZyme repertoire of a lichen fungal symbiont genome comes from a mycorrhizal study by Miyauchi *et al.*^48^ that included a single lichen, *Usnea florida,* assessed as an “endophytic lifestyle”. This lichen exhibited a profile similar to that of ectomycorrhizal fungi, sharing a relatively low enzymatic count of PCWDEs but with a considerably smaller genome size, closer to that of other endophytic fungi^48^. A second attempt to characterize lichen genomic profiles, which included 44 genomes from a single taxonomic class, found similar genomic signatures^15^. However, this time Resl *et al.*^15^ identified a secondary profile in lichen symbioses within the *Ostropomycetideae* (*Lecanoromycetes*), revealing that they retain enzymatic machinery capable of degrading plant cell walls. The proven activity of these enzymes suggests the retention of additional mechanisms for carbon acquisition beyond the commonly described nutritional symbiosis. Shortly after, a third attempt by Song *et al*.^53^ included six lichens fungal genomes, from two different taxonomic classes— *Lecanoromycetes* and *Eurotiomycetes*. However, this time all six genomes showed a reduced enzymatic profile with notable losses in the PCWDEs and sugar transporters. These reductions were mainly attributed to the close interaction with algal cells, whose cell-wall composition lacks pectin contrary to those of other land plants. As a result, Song *et al*.^53^ suggest that PCWDEs—mainly pectin-related—became non-functional since the fungi no longer require them to ‘remodel’ or interact with algal cells. Additionally, the reliance of fungi on symbiotic-originated polyols may have rendered several sugar transporters dispensable.

Our results showed that there is more than one genomic evolutionary trend in lichen symbioses. Using a much larger sample of lichen fungal genomes than in previous studies, we found both reductions and groups of lichen fungi with massive enzymatic machinery, with PCWDE numbers comparable to those of wood-decaying fungi. We identified large genomes, similar in size to those of ectomycorrhizal fungi, alongside some of the smallest genomes within the filamentous fungi in the class Ascomycota. However, all these genomes exhibited fewer annotated genes than the average of other ascomycotan genomes—including CAZymes, proteases, and PFAMs. These multiple genomic profiles mirror what has been previously observed in mycorrhizal symbioses, suggesting that photobiont carbon subsidies may serve a nutritional role in some, but not all, lichen symbioses.

### Could phylogenetic relatedness explain differences in CAZyme richness?

Our results clearly demonstrate the presence of more than one genomic CAZyme profile. Is this variation driven by different symbiotic interactions, or does it simply reflect conservatism among evolutionary clades? In Resl *et al*.^15^, the sole reported case of a divergent genomic signature occurs within a single clade. However, this clade shares more than just an evolutionary history; it also includes saprotrophs, borderline lichen symbioses—lichens with no clearly defined interaction but growing near algae—and trentepohlian lichens. This suggests that additional, unaccounted-for variables may be influencing this trend.

In a different scenario, the taxonomic class *Lichinomycetes*—which has been previously studied due to its small genome size in both lichen^27^ and non-lichen^60,61^ species—exhibits uniformly small genomic profiles, considerably smaller than those of many other lichen symbioses. This genomic signature persists despite the clade including lichen symbioses with trentepohlian-, trebouxiophycean-, and cyanobacteria-associated photobionts, and it remains indistinguishable from their closely related, non-lichenized relatives engaged in different symbiotic lifestyles—including the aforementioned ericoid mycorrhizae. These patterns suggest that the observed genomic signature may have been inherited from ancestral clade-level events, which appears to have limited the acquisition of new genomic content rather than being shaped by symbiotic association.

This conundrum is particularly evident in our results for the genomes of the classes *Dothideomycetes* and *Lichinomycetes*. In the case of the *Lichinomycetes* genomes, we consistently recover lichinomycete genomic profiles among the groups with the smallest gene sets, regardless of their associated photobionts or subsidies (Table S6), supporting a clade-driven genomic signature^27^. Although we found differences between lichen and non-lichenized *Lichinomycetes*, these differences were limited to PFAM and CAZyme annotations (Table S7). In *Dothideomycetes*—a class that includes some of the largest enzymatic repertoires among lichen symbioses—all lichens analyzed in this study belong to a single clade, the order *Trypetheliales*. Moreover, all known lichen symbioses within this class involve the same group of trentepohlian photobionts, making it challenging to disentangle the effects of subsidies, photobiont identity, or clade-specific genomic signatures. Nevertheless, our results reveal a uniform genomic profile within *Dothidiomycetes*.

A strength of the present analysis is that it compares multiple independent origins, subsidies and photobiont interaction across the ascomycotan tree of life. When we analyze our data collectively—even with the aforementioned disparate fungal genomes— a distinctive pattern emerges and the phylogenetic conservatism of nested clades becomes less obvious. Instead, the convergence of multiple independent groups within distinct genomic profiles emerges as the common denominator. This can be seen among the largest enzymatic repertoires, a genomic profile shared by species from four independent classes. Conversely, the smallest enzymatic repertoires, found in species from two classes—*Lichinomycetes* and *Lecanoromycetes*—encompass both the smallest and largest genomes known in lichen symbioses. While a shared evolutionary history may play a role in the long-term establishment of these CAZyme profiles, results suggest that it is neither the sole factor nor the primary driver of genome evolution.

### Is the subsidy the ultimate CAZyme profile driver?

The idea that algae provide their fungal symbionts with ‘assimilates’ has been around since the first description of lichen symbiosis^4^. We now understand that these assimilates encompass a variety of carbon molecules specific to each photobiont^1^. In green algal photobionts, these primarily include acyclic polyols—namely, erythritol, ribitol and sorbitol— whereas in cyanobacteria it is primarily glucose and the non-reducing disaccharide trehalose. Experimental work conducted in the 1960s has shown that the photobionts are producing these photo-assimilates and releasing them into the symbiosis inter-space^14^. In some cases, it has been reported that the symbiosis drives the production and accumulation of these compounds, with some isolates of photobionts— *Coccomyxa, Hyalococcus* and *Nostoc*— ceasing production entirely after a few hours outside the symbiosis^62^. Today, we know that some of these non-reducing sugars can also play non-nutritional roles in algae, as they are produced outside the symbiosis as an osmotic response and are rather common^34,35,36^.

In fungi, non-reducing sugars—specifically polyols—constitute the primary component of their soluble intra-cellular carbohydrates^12^. The six-carbon mannitol is the most abundant polyol by dry weight in filamentous fungi^42^. Mannitol has been largely studied in plants, algae and fungi, due to its versatility as a storage molecule, osmo-protectant and antioxidant^63^. Lichen fungi also contain relatively high levels of mannitol, followed by the five-carbon polyol arabitol; and in some scenarios constituents include volemitol, erythritol and xylitol^64^.

In the context of the symbiotic interaction, the polyols produced by green algae and the glucose derived from the cyanobacteria are released into the shared extracellular polysaccharide matrix of the symbiosis^64^. Active transporters facilitate the movement of glucose, whereas polyols rely largely on passive diffusion^65^. Ribitol has been experimentally traced by isotope-labelled NaHC^14^O3—moving from algal cells to fungal cells and assimilated into two main polyol pools: arabitol and mannitol^14,16,66^. The process is not fully understood, but a hypothesis from the 1960s suggests that ribitol is initially oxidized to ribulose and then reduced to arabitol via an NADP-dependent arabitol dehydrogenase^12,14^. A portion of this arabitol gets incorporated into the pentose phosphate pathway as a precursor of fructose, which subsequently enters glycolysis. Mannitol—well known in fungal physiology—pools via glycolysis by the reduction and later dephosphorylation of fructose-6-phosphate—F6P^12,14,63^.

In glucose-subsidized lichen symbioses, glucose—whose metabolic fate parallels that of hexoses in mycorrhizal associations—is thought to be primarily integrated into glycolysis and ultimately accumulates as mannitol following F6P dephosphorylation^16^. Sorbitol, only known from the algal order *Prasiolales*, is converted into the seven-carbon volemitol polyol by the fungi—an assimilation restricted to *Eurotiomycetes—*which in non-lichenized fungi experiments has growth curves analogous to mannitol and glucose^67^.

One theme shared across these ‘assimilates’ is that their incorporation into fungal metabolism occurs without a starting delay; suggesting that the enzymes required are central and readily available, even if sometimes they operate at a relatively slow pace.

Erythritol metabolism is a different story. In fungal cells, erythritol is typically localized in the cytoplasm, and its production has been largely attributed to an osmotic stress response, a phenomenon observed from yeast to filamentous fungi across the kingdom^68^. The production is derived from the intermediate erythrose-4-phosphate in the pentose phosphate pathway; however, although fungi can catabolize erythritol, experimental evidence indicates that it is neither preferred nor central to their primary metabolism^69^. This was inferred primarily due to a significant delay in its utilization, suggesting that its catabolism is induced by starvation, and it is dependent on a NAD-dependent polyol dehydrogenase with high substrate specificity^70,71^. More recently, the genes involved have been identified in the yeast *Lipomyces* and *Yarrowia,* where an erythritol dehydrogenase and erythrulose kinase were characterized; subsequent knockout experiments confirmed their roles, as mutants exhibited impaired growth on erythritol-enriched media^72,73^.Although the complete erythritol cycle remains incompletely understood in fungi, it is thought that erythritol could be reincorporated into glycolysis via a hypothetical catabolic pathway of erythrulose-phosphate as a probable precursor of dihydroxyacetone phosphate (DHAP) and formaldehyde, analogous to erythritol usage in bacteria^72^. More recent studies include a recycling step where erythrulose-1-phosphate is converted to erythritol-4-phosphate via an isomerase, as the last step in erythritol catabolism^74^.

In erythritol-subsidized lichen symbioses, the erythritol detected is a by-product of the trentepohlian algae—or the trebouxioid genus *Apatococcus*^34^—photosynthesis, not the fungi^16,33^. Erythritol fate has been traced using isotope labelled NaHC^14^O3 which gets assimilated via algal photosynthesis and later released to the symbiosis. However, unlike other photo-assimilates, erythritol is subsequently washed out with little (<2∼9%) to no signs of fungal assimilation^14^. Contrary to other ‘assimilates’, erythritol seems not to serve a nutritional function and it has been hypothesized to act as a “physiological buffering” or osmo-protectant through rewetting or drying cycles^16,18^. This functionality has also been proposed to explain the dual nature of polyol pools in the ribitol-subsidized lichen symbioses, where one of the polyols functions as a desiccation protectant, and the other is a readily available carbon source for fungal respiration^1^. In erythritol-subsidized lichen symbioses—although in some cases it has been reported to occur together with arabitols and traces of ribitol^33^—this osmolyte role cannot be maintained unless there is another carbon source available to the fungi.

Our results showed a differential enzymatic machinery between the erythritol-subsidized fungal genomes compared to genomes with other subsidy molecules. However, this change in machinery is not required nor involved in erythritol usage or incorporation into glycolysis or the glycolytic pathway. The enzymatic arsenals here described are mainly related with plant cell wall degradation suggesting that the machinery is still required by the fungi either to interact with the cell walls of the photobiont—softening or remodelling to facilitate exchanges—or to harvest carbon from other sources.

### Does photobiont cell wall composition shape the profile of retained CAZymes?

As mentioned by Song et al.^53^, algal cells have different cell wall composition, a factor that may have driven a differential enzymatic reduction in genomes. Considering that photobionts range from bacteria, heterokonts and include representatives from three different taxonomic classes of green algae—as far as we know—, the picture becomes even more complex, with not a sole outcome. In this study we included representatives from two taxonomic classes of green algae together with cyanobacteria, thereby covering a relatively broad spectrum of cell wall composition.

Cyanobacterial cell walls in lichens are no different than their free-living counterparts or other bacteria; their main structural constituent is peptidoglycan^76^. In addition, the cells come surrounded by an extracellular polysaccharide matrix that can be distinguishable from fungal polysaccharides by their additional complexity, which usually come with more than six monosaccharides, with a high number of glucose residues and often incorporate D-glucosyluronic acid and D-xylopyranose residues^77^. These features are also standard in free-living forms, with some specific extracellular polysaccharides occurring in their free-living forms but not within lichen symbioses. Our results show that the genomes of lichen fungi associated with cyanobacteria experienced the highest reduction in their enzymatic arsenals, including most PCWDEs (except for AA9), and the complete absence of the peptidoglycan recycling family CE9, which was present across all other genomes— suggesting that no specific enzymes are required to improve carbon accessibility, and potentially reflecting a nutritional symbiotic relationship.

Green algae, by contrast, exhibit an entirely different suite of cell wall constituents. For instance, *Trebouxiophyceae* algae are known to have multilayered polysaccharide-based cell walls, with main constituents collectively including cellulose, glucomannans, xylans, algaenans and β-galactofuranan^78,79,80^. However, some orders of algae involved as lichen photobionts—such as *Chlorelalles*, *Trebouxiales* and *Coccomyxa—*have recurrently shown either a complete lack of cellulose and pectin^79^ or, in the case of *Trebouxia* and *Pseudotrebouxia* (now *Trebouxia*), trace amounts of cellulose associated with inner cell wall layers^81^. Despite these differences, at the genus level *Trebouxia* and *Asterochloris* cell walls are predominantly composed of β-xylorhamnogalactofuranan units^82,83^. In the order *Prasiolales* the free-living algae *Prasiola* has been repeatedly described as having an inner cellulose layer^84^, while the lichen symbionts *Stichococcus* and *Diplosphaera* have only been indirectly assessed based on similarities to *Chlorella* and *Chloroidium* from the order *Chlorelalles*^85^.

The overall smaller number of in PCWDEs in lichen fungal genomes compared to non-lichen fungi—evidenced by the near absence of polysaccharide lyases (PL), a significant decrease of AA9 cellulases, and the complete lack of the complementary cellulose-binding domain (CBM1)—suggests a reduced interaction with the plant cell wall, whether in carbon harvesting or cell wall remodeling. However, if these trait losses were solely a consequence of the lower cellulose content in the algal cell wall, one would expect to see an increase in compensatory traits related to the differential composition of the specific cell walls. In the case of *Trebouxiales*, the only CAZyme family that exhibited collectively a significantly higher number of annotations—when compared against other lichen symbioses—was the GH128 family of glucan endo-1,3-β-D-glucosidases, which hydrolyze glycosidic bonds in glucans. Still, it is not specific to the cell wall. Additionally, when examining the genomes of sorbitol– and ribitol-subsidized fungi together—*i.e*. *Trebouxiales* and *Prasiolales*—, we observed that the GT25 family of galactosyltransferases—responsible for catalyzing the transfer of galactose—doubled in occurrence (from 1 to 2 annotations) relative to cyanobacterial– and trentepohlioid-lichen fungal genomes. However, none of the significantly increased enzymatic annotations can be directly linked to either β-xylorhamnogalactofuranan or β-galactofuranan activity— suggesting that our data do not exhibit a pattern of trait compensation that can be explained solely by differences in cell wall composition.

In the case of Ulvophyceaen algae, there are fewer comprehensive studies behind the filamentous *Trentepohliales*, which includes the lichen photobionts: *Cephaleuros*, *Printzina*, *Phycopeltis* and *Trentepohlia*. Existing research has primarily relied on microscopy and stain-based methods to characterize the cell wall layers. The genus *Trentepohlia* has been reported to contain both cellulose and pectin, with a ‘pectose’ cap in the apical growth region; among these, cellulose is the most frequently reported^86,87,88,89^. In contrast, stain-based analysis of the genus *Phycopeltis* identified sporopollenin to be the most common constituent, while cellulose was not detected—a possible cellulose masking phenomenon, that the same authors suggest may be attributed to the presence of sporopollenin^90^. Similar considerations were later reported in *Trentepohlia*^91^. Additionally, in the lichen genus *Roccella* (*Arthoniomycetes*) with a trentepohlian-photobiont (*Trentepohlia*), cellulose was not found in a whole-thallus analysis. The glycosyl linkage analysis did not report 4-linked glucose residues (4-Glc)— the gold standard for cellulose—and the glucose detected was related to fungal-derived galactomannans, which are the dominant cell wall components in the lichen thallus^92^.

The duality of the plant cell wall composition in *Trentepohliales—*based on the literature*—* seems similar to the bimodal distribution of our annotated enzymatic profiles. We observed two distinct groups of trentepohlioid-lichen genomes: one comprises a substantial number of annotated enzymes putatively targeting the canonical trentepohlian cell walls components, while the other group carries CAZyme repertoires closer to those of trebouxiophycean-lichens, with a markedly fewer number of PCWDEs. This enzymatic dichotomy also echoes the two phylogenetic clades of lichen-derived trentepohlioids described by Nelsen *et al*.^93^, though we do not know if these possess different cell wall composition. Those clades comprised intermixed species from the *Trentepohlia* and *Printzina*^94^, and at the same time they associated with lichen fungal symbionts from different taxonomic classes, yet their split remains unexplained. Here, we treat photobionts from both genera simply as trentepohlioids, without differentiating them due to their unresolved taxonomic and phylogenetic placement^95,96^, and like Nelsen et al. — with algae—we found no distinctive genus-specific clustering within their lichen fungal symbionts. Notably, our dataset contains lichens closely related to those analysed in Nelsen et al., and our bimodal enzymatic pattern aligns with their phylogenetic clades, suggesting that there could be an intrinsic biological difference between trentepohlioid algae that may be differentially shaping their lichen fungal partners.

### What can we say, and what we cannot

One of our goals was to step away from the published records and read through the lichen fungal genomes, looking for symbiotic evolution signatures that could occur concurrently with their symbionts, subsidies, or close relatives. Lichen research continues to show progress, and nowadays there are reported cases of multiple algae coexisting in a single thallus—a condition previously known only from some species. Still, it has recently become a common feature with the incorporation of metabarcoding analysis, and some of these co-occurring algae have been observed under the microscope within the same thallus (*Trebouxia* and *Coccomyxa* within a thallus of *Protoparmelia muralis*)^97,98^. This promiscuity was not accounted for in our surveys. In some scenarios, considering that many algae within the trebouxioid group share the same polyols and cell wall compositions, this promiscuous behaviour makes sense given the here-exposed enzymatic profiles, but ultimately limits the range of our inferences.

Another example of our current limitations is that lichen symbiosis studies primarily focus on fungi that partner with members of the *Trebouxiales*. This is not a consequence of researcher bias towards specific groups, but rather the current diversity distribution of the lichen symbioses, where *Trebouxia* is one of the most common photobionts of lecanoromycete lichens. However, some photobionts are more widely spread across lichen symbioses than others, with some fungal classes only in symbioses with photobionts different from *Trebouxia* — e.g. *Dothideomycetes*, *Eurotiomycetes*, and *Leotiomycetes*.

Our results support the notion that some lichen fungal genomes resemble non-nutritional symbiotic profiles within erythritol-subsidized genomes, while others seem to have reduced tools to harvest external carbon sources. We can, therefore, circumscribe two evolution paths irrespective of their fungal taxonomic classes. However, we are limited in whom to attribute these effects, whether to the algal cell walls or the subsidies, including the still intractable distinction within *Trentepohlia* sensu lato. To unambiguously attribute the shaping of genomic carbohydrate-degrading machineries, we will require better characterizations of the algal cell-wall composition, together with *in vivo* demonstrations of the enzyme activity action in lichen symbiosis.

## Materials and methods

### Sample selection and data curation

We surveyed the NCBI Assemblies and Reference Sequences database for all lichen-derived genomes, either from axenic cultures or metagenome-assembled genomes, up to February 2024. To ensure a comprehensive coverage of all available data, we manually browsed the NCBI Assembly and Genome database and filtered across all known taxonomic orders that encompass lichen symbioses within *Ascomycota*^22^. These included taxa labelled as *incertae sedis* at phylum, class, and order levels (Table S1).

Our search in the NCBI Assemblies database yielded 189 entries, representing 130 species across 85 genera. Due to the expected variation at the genus level—*e.g.* photobionts, morphology, genomics —, we chose to retain all available species for each genus to capture the full extent of genomic diversity. We opted to use one representative genome per species, with carefully considered exceptions. These exceptions were retained based on the rationale of our statistical analyses, which employ global estimates that would benefit from additional genomic diversity without introducing bias. For example, we included three genomes from the lichen fungi *Endocarpon pusillum* from the class *Eurotiomycetes*, which associates with a photobiont from the *Prasiolales* clade, which is scarcely represented in our dataset. We also retained genomes from underrepresented fungal taxonomic classes, such as *Bathelium mastoideum* (2) and *Viridothelium virens* (2) in the class *Dothidiomycetes*. Additionally, we included one genome from the subclass *Lecanoromycetidae sp.*, which lacks species or photobiont identification, and the wrongly identified *Pseudosagedia rhaphidosperma* as established by the revision of Tagirdzhanova et al. (2024). Although these genomes were included in global lichen fungal statistics, it was excluded from analyses focusing on photobiont associations and subsidy molecule statistics. Finally, we added the external genome assembly of *Graphis scripta* published by Resl *et al.*^15^ but not currently included in the NCBI Assemblies. A total of 130 genomes were kept after filtering (Table S1).

Using our NCBI survey as a baseline, we collected all available photobiont information from the literature up to the genus level (Table S3). Then, we decided to expand our genomic survey to increase the yield from the underrepresented photobiont associations—mainly cyanobacterian– and trentepohlian-lichen symbioses. We used the lichen metagenomes libraries database from Tagirdzhanova *et al.*^99^, which includes all lichen-derived NCBI Sequence Read Archive (SRA) libraries from non-culture-based studies, and we filtered out those already represented in the available assemblies. We retained all libraries for cyanobacterian and trentepohlian lichens (15 and 20, respectively), as well as libraries from the Ascomycota class *Eurotiomycetes*, to increase the samples with photobionts from the *Prasiolales* order. The resulting 38 libraries selected from the SRA included 38 species from 29 genera.

We further sought to include underrepresented class-level fungal groups and photobiont to generate new genomic data. This effort resulted in the inclusion of samples from *Arthoniomycetes* (six—five from lichen symbioses and one saprotroph), *Leotiomycetes* (1), *Lecanoromycetes* (12), *Eurotiomycetes* (1), and species with an unassigned taxonomic class (4). In total, we selected 24 new samples for genome sequencing, including 23 lichens distributed among 22 species from 21 genera. These samples comprise the non-lichenized *Arthonia sanguinea*, one undescribed lichen species and two samples from the *incertae sedis* species *Wadeana dendrographa*.

In total, our study included 191 lichen fungal genomes (Table S2). To provide a contrast to these lichen-derived fungal genomes, we incorporated 118 non-lichen fungal genomes, representing all classes within filamentous Ascomycota. Of these, 117 genomes were sourced from the assembly sets published by Resl *et al.*^15^ and Díaz-Escandon *et al.*^27^, supplemented by the newly sequenced *Arthonia sanguinea* from the *Arthoniomycetes*, and the saprotrophic *Mycocallicium subtile* from the *Eurotiomycetes*, which was obtained from the NCBI SRA database. In sum, 309 genomes were used in this study.

### DNA extraction and sequencing

We used a variety of fresh and old material (Table S2). We aimed for ascocarps to increase fungal cell density in the scenarios where it was feasible, together with the thallus to get algal DNA whenever it was possible. All the DNA extraction steps were conducted at the University of Alberta. Samples were frozen at –80°C overnight and grinded using Qiagen TissueLyser II (Qiagen, Germany) with tungsten beads at 25Hz for 30 seconds or until pulverized. We used the QIAamp DNA investigator kit (Qiagen, Germany) for DNA extraction following manufacturer’s protocol, with an overnight lysis step and two elution rounds with 40uL and 30 minutes incubation periods. DNA quality was assessed using Implen NP80 nanophotometer (Implen, Germany) and quantified using Invitrogen Qubit 2.0 fluorometer (Life Technologies/Thermofischer, Waltham, MA, US). For the short-read sequencing, we used Genome Québec (Canada) as an external service provider for metagenomic library preparation and sequencing. The sequencing was performed using NovaSeq 6000 PE 150 bp technology (Illumina, San Diego, CA, US).

### Metagenomes assemble and genomic binning

Newly generated metagenomic libraries, along with SRA data, were assessed for quality using FastQC (v0.11.9)^100^. Sequences were trimmed and filtered using Trimmomatic to remove adapters (TruSeq3-PE-2) and reads with a quality score below Q30. The metagenomes were assembled using metaspades^101^ in paired-end reads mode with *k*-mers of 21, 31, 51, 71, 81, 101, and 127. The resulting assemblies were binned using CONCOCT (v1.1)^102^, and each bin was assessed for eukaryotic content using Eukcc (v0.1.5.1)^103^. The eukaryotic-passed bins were tested with BUSCO (v5.2.3) for completeness and contamination^104^.

Subsequently, all bins (both passed and unpassed) were manually inspected using the ‘GC content vs Coverage’ plot method from Tagirdzhanova *et al*.^105^ to check for inconsistencies and potential errors during the binning process. This step allowed us to merge bins that may have been erroneously split, despite their similarity, by carefully inspecting for duplication accumulation in BUSCO and cross-validating gene content in the merged bins. After merging, we inspect all bins for completeness (Table S1). The refined bins were taxonomically assessed using nuclear gene markers to verify the species from the library source. The genome *Pseudosagedia rhaphidosperma*, previously reported as a misidentification^99^, was confirmed and belonged to *Teloschistales*. We retained it for class-level analysis but later excluded it from the photobiont and subsidy analyses.

### Gene prediction and functional annotation

All the Metagenome Assembled Genomes (MAGs) were cleaned, masked, predicted and functionally annotated using the Funannotate pipeline (v1.8.15)^106^. For the cleaning process, we applied the ‘--exhaustive’ option to process every contig for duplicate removal, and then we sorted the contigs without filtering at any length. For masking, we set the run to use the default masking tool TANTAN^107^. Gene prediction was performed using GeneMark-ES in the fungal mode (--fungus) and combined in the funannotate prediction pipeline using the option ‘--genemark_gtf’. Funnanotate first carries out an initial BUSCO prediction using a modified version of BUSCO2 with the DIKARYA_ODB9 dataset and an Augustus reference (--species=anidulans). These validated gene models are then used to train an Augustus (v3.3.2)^108^ model and perform an *ab initio* prediction with a subsequent optimization step (--optimize_augustus), followed by gene predictions using SNAP (v2006-07-28)^109^ and glimmerHMM (v3.0.4)^110^. Finally, funannotate prediction produces a consensus gene structure annotation using EVidenceModeler (v.1.1.1)^111^, together with tRNA predictions generated by tRNAscan-SE (v2.0.9)^112^.

For functionally annotations we used funannotate together with InterProScan5 (v5.63-95.0)^113^ and eggNOG-mapper (v 2.1.12)^114^, using eggNOG database (v5.0.2)^115^. We then integrated all annotation using the Funannotate annotate step, which employs PFAM (v36.0)^116^ and Uniprot (v2023_05)^117^ for protein families and function annotations, dbCAN HMMdb (v12.0)^118^ for carbohydrate-active enzymes (CAZymes), and MEROPS (v12.0)^119^ for proteases annotations.

For genomes derived from the NCBI Assemblies and Reference database, we initially inspected those previously published that used funannotate for prediction and annotation. Genomes without previous annotations were predicted and annotated following the steps described above. In the case of genomes previously annotated and predicted with funannotate, we downloaded their annotations from NCBI in genbank format and run the ‘annotate’ step from funannotate pipeline to update the annotations with the most recent databases (PFAM v36.0, Merops v12.0, dbCAN v12.0, Uniprot v2023_05). Additionally, some publications have included their GFF3 annotations together with their proteins, but they are not available in NCBI, in these scenarios we combine them using the ‘annotate’ step to reconcile the data and update annotations. This guarantees that all the genomes here analyzed were annotated with the same databases, regardless of the source. We limit this step to known funnanotate predictions only (Table S1).

### Statistical analyses

We combine all the annotation files produced by funannotate using custom Python scripts (github). The statistical analyses were conducted using R (v4.4.1), and we compared global metrics including genome size, number of genes, GC content, number of tRNAs, total annotated PFAMs, CAZymes and proteases. Custom scripts were used to determine whether to apply Wilcoxon rank-sum test or T-test for each dataset, based on assumptions underlying each method. In addition, we produced density plots to inspect the distribution of each dataset and to visually contrast these distributions with the test results. Based on the observed different multi-peak density distributions, we tested for multimodality, using two approaches: Hartigan’s Dip Test (included in package ‘dip.test’) which assumes a single mode as the null hypothesis, and Silvermman’s test (included in the package ‘multimod’) which allows testing for *k* modes (mod0) using permutations.

To further investigate the difference between lichen fungal symbionts and non-lichen fungi, we group the genomes by photobiont type and subsidy molecules, focusing on the total number of CAZyme annotations. We used the nonparametric Kruskal-Wallis test to assess differences between groups and performed a post-hoc Dunn’s test for multiple comparisons and level assignments. Additionally, we evaluated multimodality within each photobiont and subsidy group, and we calculated the proportion of CAZYme classes contributing to each subsidy.

To further analyse CAZymes differences among the subsidy molecules, we filtered the dataset to include only lichen fungal genomes. Using a custom script, we evaluated differences across subsidies by iterating over each CAZyme family individually. The script employs a two-steps approach: first, it runs a nonparametric Kruskal-Wallis (KW) test, and then it fits Generalized Linear Models (GLMs) using either Poisson or negative binomial distribution, depending on the level of overdispersion (dispersion parameter < 1.5 for Poisson models) and based on each model’s Akaike Information Criterion (AIC). For the GLMs, the resulting models for each CAZyme family were compared against a null model using a Likelihood-Ratio Test (LRT) to assess whether the subsidy variable significantly explains the observed differences. In both tests we corrected the *p* value using the Bonferroni correction and the Benjamini-Hochberg (BH) procedure. For pairwise post hoc analyses, we used the nonparametric Dunn’s test for KW-test results and Estimated Marginal Means (EMMs) for the GLMs results. All instances of post hoc tests used the BH correction.

To validate these comparisons, we did not filter the post hoc analyses solely based on the global p-value. Instead, since we ran two independent tests to assess significance, we kept the results together for manual inspection and noted cases where both tests reached a consensus, regardless of their post-hoc arrangements. All p-values were adjusted using the Bonferroni correction and the Benjamini-Hochberg (BH) method, prioritizing the post hoc contrasts with “BH” to balance the false discovery rate and maintain statistical power. Finally, we manually inspected the results against a heatmap summarizing all CAZyme families to confirm the statistical findings visually. For the latter applications, we reported the *p*-values corrected using the BH procedure and the results from Dunn’s test, as this test was less sensitive to sample size.

### Phylogenetic reconstruction

To produce a reference phylogeny for posterior clustering analyses, we implemented the Phylociraptor (v1.0)^120^ pipeline, including all the predicted proteomes from our functional annotation step. We set the pipeline to use the ‘ascomycota_odb10’ BUSCO set (1706) and used BUSCO (v5.2.4)^104^ to collect all matching proteins per proteome. Then, we filtered proteomes based on a minimum of 20% completeness and kept only proteins occurring in 30 or more proteomes. The resulting 1387 genes were aligned using mafft (v7.464)^121^ with automatic optimization (--auto) and trimmed using Clipkit (v2.3.0)^122^ with default settings. The alignments were filtered with a threshold of 50 parsimony sites or more in and ran model testing using IQTREE2(v2.0.7)^123^. We produced gene trees for each alignment set and using the mean bootstrap of the gene trees we filtered up genes with mean bootstrap above 80%. Finally, we produce a species tree using ASTRAL(v5.7.1)^124^.

### Clustering analyses

To evaluate the CAZyme family composition across all genomes and along subsidies, we computed a scaled (*z*) principal component analysis (PCA) using only the lichen fungal genomes and the function ‘prcomp’ in R ‘stats’. In addition, given that our CAZyme model consisted of over 162 parameters—and some of these could be correlated at different taxonomic levels, thereby oversplitting variance—we test for phylogenetic signals across the first two axes. We use the previously constructed phylogeny, and we calculate Pagel’s λ using ‘phylosig’ implemented in the R package ‘phytools’^125^ Additionally, we computed a phylogenetically corrected PCA (p-PCA), using the phylogenomic tree produced above and the function ‘phyl.pca’ from the R package ‘phytools’^125^. Finally, we evaluated the contribution to the clustering of each CAZyme class using the median of loading weights across CAZyme families per component in both the PCA and pPCA, and scaled these values by the maximum coordinates per axis in the final plots.

## Data availability

All newly generated genomes, including libraries and annotated metagenomically assembled genomes, are available at the National Center for Biotechnology Information (NCBI) under BioProject PRJNA1472413. Additionally, code and protein predictions produced from externally produced genomes are available at Figshare, doi: 10.6084/m9.figshare.33106691. Intermediate files that exceed the size limit allowed by Figshare can be obtained upon request to the mailing author.

## Acknowledgements

This research was funded by an Alberta Graduate Excellence Scholarship and Doctoral Colombian Minciencias grant to DDE; a Natural Sciences and Engineering Research Council of Canada (NSERC) Discovery Grant (RGPIN-2019-04892) to TS; and a Canada Research Chair in Symbiosis to TS. Data analyses associated with this study were performed with support from the Digital Research Alliance of Canada (formerly Compute Canada).

## References

1. Spribille T, Resl P, Stanton DE, Tagirdzhanova G. Evolutionary biology of lichen symbioses. New Phytol.: 2022; 234:1566–1582. 10.1111/nph.18048

2. Honegger R. Functional aspects of the lichen symbiosis. Annu Rev Plant Physiol Plant Mol Biol. 1991;42(1):553–7. 10.1146/annurev.pp.42.060191.003005

3. Pichler G, Muggia L, Carniel FC, Grube M, Kranner I. How to build a lichen: from metabolite release to symbiotic interplay. New Phytol. 2023 May 4;238(4):1362–78. doi:10.1111/nph.18780

4. Schwendener S. Die Algentypen der Flechtengonidien. Basel: Universitaetsbuchdruckerei; 1869.

5. Lücking R, Spribille T. The Lives of Lichens: A Natural History. Princeton University Press; 2024.

6. Harley JL, Smith DC. Sugar Absorption and Surface Carbohydrase Activity of Peltigera polydactyla (Neck.) Hoffm. Ann Bot. 1956;20(80):513–43.

7. Smith DC. Studies in the physiology of lichens: Experiments with dissected discs of *Peltigera polydactyla*. Ann Bot. 1960;24(2):186–99. doi:10.1093/oxfordjournals.aob.a083694

8. Smith DC. Studies in the physiology of lichens: IV: carbohydrates in *Peltigera polydactyla* and the utilization of absorbed glucose. New Phytol. 1963;62(2):205–16. doi:10.1111/j.1469-8137.1963.tb06327.x

9. Bednar TW, Smith DC. VI. Preliminary studies of photosynthesis and carbohydrate metabolism of the lichen xanthoria aureola. New Phytol. 1966;65(2):211–20. doi:10.1111/j.1469-8137.1966.tb06353.x

10. Drew EA, Smith DC. The Physiology of the Symbiosis in *Peltigera polydactyla* (Neck.) Hoffm. Lichenol. 1966;3(2):197–201.

11. Richardson DHS, Smith DC. The physiology of the symbiosis in *Xanthoria aureola* (ach.) erichs. Lichenol. 1966;3(2):202–6. doi:10.1017/S0024282966000215

12. Lewis DH, Smith DC. Sugar Alcohols (Polyols) in Fungi and Green Plants. I. Distribution, Physiology and Metabolism. New Phytol. 1967;66(2):143–84 doi:10.1111/j.1469-8137.1967.tb05997.x

13. Richardson DHS, Smith DC, Lewis DH. Carbohydrate movement between the symbionts of lichens. Nature. 1967;214(5091):879–82. doi:10.1038/214879a0

14. Richardson DHS, Hill DJ, Smith DC. Lichen physiology: XI. The role of the alga in determining the pattern of carbohydrate movement between lichen symbionts. New phytol. 1968;67(3):469–86. doi:10.1111/j.1469-8137.1968.tb05476.x

15. Resl P, Bujold AR, Tagirdzhanova G, Meidl P, Freire Rallo S, Kono M, et al. Large differences in carbohydrate degradation and transport potential among lichen fungal symbionts. Nat Commun. 2022 May;13(1): 2634. doi:10.1038/s41467-022-30218-6

16. Farrar JF. Ecological physiology of the lichen *Hypogymnia physodes* II. Effects of wetting and drying cycles and the concept of ‘physiological buffering.’ New Phytol. 1976;77(1):105–13. doi:10.1111/j.1469-8137.1976.tb01504.x

17. Cowan DA, Green TGA, Wilson AT. Lichen metabolism. 1. The use of tritium labelled water in studies of anhydrobiotic metabolism in Ramalina celastri and Peltigera polydactyla. New Phytol. 1979;82(2):489–503. doi:10.1111/j.1469-8137.1979.tb02676.x

18. Smith D. Is a Lichen a Good Model of Biological Interactions in Nutrient-Limited Environments? In: Shilo M. Strategies of microbial life in extreme environments: report of the Dahlem Workshop on Strategy of Life in Extreme Environments. Weinheim, Germany: Verlag Chemie;1979:291–301

19. Aubert S, Juge C, Boisson AM, Gout E, Bligny R. Metabolic processes sustaining the reviviscence of lichen *Xanthoria elegans* (Link) in high mountain environments. Planta. 2007 Oct;226(5):1287–97. doi:10.1007/s00425-007-0563-6

20. Kosugi M, Miyake H, Yamakawa H, Shibata Y, Miyazawa A, Sugimura T, et al. Arabitol provided by lichenous fungi enhances ability to dissipate excess light energy in a symbiotic green alga under desiccation. Plant Cell Physiol. 2013 Aug 1;54(8):1316–25. doi: 10.1093/pcp/pct079

21. Gargas A, DePriest PT, Grube M, Tehler A. Multiple origins of lichen symbioses in fungi suggested by SSU rDNA phylogeny. Science. 1995 Jun 9;268(5216):1492–5. doi:10.1126/science.7770775.

22. Lücking R, Hodkinson BP, Leavitt SD. The 2016 classification of lichenized fungi in the Ascomycota and Basidiomycota – approaching one thousand genera. Bryologist. 2016;119(4): 361–416. doi:10.1639/0007-2745-119.4.361

23. Prieto M, Schultz M, Olariaga I, Wedin M. Lichinodium is a new lichenized lineage in the Leotiomycetes. Fungal Divers. 2019; 94: 23–39. doi:10.1007/s13225-018-0417-5

24. Wijayawardene, N.N., Hyde, K.D., Mikhailov, K.V. et al. Classes and phyla of the kingdom Fungi. Fungal Diversity 128, 1–165 (2024). doi:10.1007/s13225-024-00540-z

25. Lutzoni, F., Pagel, M. & Reeb, V. Major fungal lineages are derived from lichen symbiotic ancestors. Nature 411, 937–940 (2001). doi:10.1038/35082053

26. Schoch CL, Sung GH, López-Giráldez F, Townsend JP, Miadlikowska J, Hofstetter V, et al. The ascomycota tree of life: A phylum-wide phylogeny clarifies the origin and evolution of fundamental reproductive and ecological traits. Syst Biol. 2009;58(2):224–39. doi:10.1093/sysbio/syp020

27. Díaz-Escandón D, Tagirdzhanova G, Vanderpool D, Allen CCG, Aptroot A, Češka O, et al. Genome-level analyses resolve an ancient lineage of symbiotic ascomycetes. Curr Biol. 2022; 32(23):5209–5218.e5. doi:10.1016/j.cub.2022.11.014

28. Thüs H, Muggia L, Pérez-Ortega S, Favero-Longo SE, Joneson S, O’Brien H, et al. Revisiting photobiont diversity in the lichen family Verrucariaceae (Ascomycota). Eur J Phycol. 2011;46(4):399–415. doi: 10.1080/09670262.2011.629788

29. Sanders WB, Masumoto H. Lichen algae: the photosynthetic partners in lichen symbioses. The Lichenologist. 2021;53(5):347–93. doi:10.1017/S0024282921000335

30. Puginier C, Libourel C, Otte J, Skaloud P, Haon M, Grisel S, et al. Phylogenomics reveals the evolutionary origins of lichenization in chlorophyte algae. Nat Commun. 2024;15(4452). doi: 10.1038/s41467-024-48787-z

31. Chrismas N, Tindall-Jones B, Jenkins H, Harley J, Bird K, Cunliffe M. Metatranscriptomics reveals diversity of symbiotic interaction and mechanisms of carbon exchange in the marine cyanolichen *Lichina pygmaea*. New Phytol. 2024;241(5):2243–57. doi: 10.1111/nph.19320

32. Kremer BP. Taxonomic implications of algal photoassimilate patterns. Br Phycol J. 1980;15(4):399–409. doi:10.1080/00071618000650401

33. Feige GB, Kremer BP. Unusual carbohydrate pattern in *Trentepohlia* species. Phytochemistry. 1980;19(8):1844–5. doi:10.1016/S0031-9422(00)83826-1

34. Gustavs L, Görs M, Karsten U. Polyol patterns in biofilm-forming aeroterrestrial green algae (Trebouxiophyceae, Chlorophyta). J Phycol. 2011;47(3):533–7. doi:10.1111/j.1529-8817.2011.00979.x

35. Hotter V, Glaser K, Hartmann A, Ganzera M, Karsten U. Polyols and UV-sunscreens in the Prasiola-clade (Trebouxiophyceae, Chlorophyta) as metabolites for stress response and chemotaxonomy. J Phycol. 2018;54(2):264–74. doi:10.1111/jpy.12619

36. Gustavs L, Eggert A, Michalik D, Karsten U. Physiological and biochemical responses of green microalgae from different habitats to osmotic and matric stress. Protoplasma. 2010;243(1):3–14. doi:10.1007/s00709-009-0060-9

37. Holzinger A, Karsten U. Desiccation stress and tolerance in green algae: consequences for ultrastructure, physiological and molecular mechanisms. Front Plant Sci. 2013;4(327) doi: 10.3389/fpls.2013.00327

38. Holzinger A, Plag N, Karsten U, Glaser K. Terrestrial Trentepohlia sp. (Ulvophyceae) from alpine and coastal collection sites show strong desiccation tolerance and broad light and temperature adaptation. Protoplasma. 2023 Nov 1;260(6):1539–53. doi: 10.1007/s00709-023-01866-2

39. Farrar JF, Smith DC. Ecological physiology of the lichen *Hypogymnia physodes* III. The importance of the rewetting phase. New Phytol. 1976;77(1):115–25. doi: 10.1111/j.1469-8137.1978.tb01604.x

40. Cooper G, Carroll GC. Ribitol as a Major Component of Water-Soluble Leachates from Lobaria oregana. Bryologist. 1978;81(4):568. doi:10.2307/3242343

41. Dudley SA, Lechowicz MJ. Losses of Polyol through Leaching in Subarctic Lichens. Plant Physiol. 1987;83(4):813–5. doi:10.1104/pp.83.4.813

42. Solomon, Peter S., Waters, Ormonde D.C., Oliver, Richard P. Decoding the mannitol enigma in filamentous fungi. Trends Microbiol. 2007;15(6):257–262. doi:10.1016/j.tim.2007.04.002

43. van der Heijden MGA, Martin FM, Selosse M, Sanders IR. Mycorrhizal ecology and evolution: the past, the present, and the future. New Phytol. 2015;205(4):1406–23. doi: 10.1111/nph.13288

44. Martin FM, van der Heijden MGA. The mycorrhizal symbiosis: research frontiers in genomics, ecology, and agricultural application. New Phytol. 2024;242(4):1486–506. doi: 10.1111/nph.19541

45. Martin F, Kohler A, Murat C, Veneault-Fourrey C, Hibbett DS. Unearthing the roots of ectomycorrhizal symbioses. Nat Rev Microbiol. 2016;14(12):760–73. doi:10.1038/nrmicro.2016.149

46. Morin E, Miyauchi S, San Clemente H, Chen ECH, Pelin A, de la Providencia I, et al. Comparative genomics of *Rhizophagus irregularis*, *R. cerebriforme*, R. diaphanus and Gigaspora rosea highlights specific genetic features in Glomeromycotina. New Phytol. 2019;222(3):1584–98. doi:10.1111/nph.15687

47. Murat C, Payen T, Noel B, Kuo A, Morin E, Chen J, et al. Pezizomycetes genomes reveal the molecular basis of ectomycorrhizal truffle lifestyle. Nat Ecol Evol. 2018;2(12):1956–65. doi: 10.1038/s41559-018-0710-4

48. Miyauchi S, Kiss E, Kuo A, Drula E, Kohler A, Sánchez-García M, et al. Large-scale genome sequencing of mycorrhizal fungi provides insights into the early evolution of symbiotic traits. Nat Commun. 2020;11(1):5125. doi:10.1038/s41467-020-18795-w

49. Martino E, Morin E, Grelet G, Kuo A, Kohler A, Daghino S, et al. Comparative genomics and transcriptomics depict ericoid mycorrhizal fungi as versatile saprotrophs and plant mutualists. New Phytol. 2018 Feb 7;217(3):1213–29. doi:10.1111/nph.14974

50. Perotto S, Daghino S, Martino E. Ericoid mycorrhizal fungi and their genomes: another side to the mycorrhizal symbiosis? New Phytol. 2018 Dec;220(4):1141–7. doi:10.1111/nph.15218

51. Chen J, Tang Y, Kohler A, Lebreton A, Xing Y, Zhou D, et al. Comparative Transcriptomics Analysis of the Symbiotic Germination of *D. officinale* (Orchidaceae) With Emphasis on Plant Cell Wall Modification and Cell Wall-Degrading Enzymes. Front Plant Sci. 2022 May;13:880600. doi: 10.3389/fpls.2022.880600

52. Gong Y, Lebreton A, Zhang F, Martin F. Role of carbohydrate-active enzymes in mycorrhizal symbioses. Essays Biochem. 2023 Apr 18;67(3):471–8. doi: 10.1042/EBC20220127

53. Song H, Kim K-T, Park S-Y, Lee G-W, Choi J, Jeon J, et al. A comparative genomic analysis of lichen-forming fungi reveals new insights into fungal lifestyles. Sci Rep. 2022 Jun 24;12(1):10724. doi:10.1038/s41598-022-14340-5

54. Lemieux C, Otis C, Turmel M. Chloroplast phylogenomic analysis resolves deep-level relationships within the green algal class Trebouxiophyceae. BMC Evol Biol. 2014;14(1):211. doi:10.1186/s12862-014-0211-2

55. Bennett GM, Moran NA. Heritable symbiosis: The advantages and perils of an evolutionary rabbit hole. Proc Natl Acad Sci U S A. 2015;112(33):10169–76. doi:10.1073/pnas.1421388112

56. Kohler A, Kuo A, Nagy LG, Morin E, Barry KW, Buscot F, et al. Convergent losses of decay mechanisms and rapid turnover of symbiosis genes in mycorrhizal mutualists. Nat Genet. 2015 Apr 23;47(4):410–5. doi:10.1038/ng.3223

57. Wu G, Miyauchi S, Morin E, Kuo A, Drula E, Varga T, et al. Evolutionary innovations through gain and loss of genes in the ectomycorrhizal Boletales. New Phytol. 2022;233(3):1383–400. doi:10.1111/nph.17858

58. Rosling A, Eshghi Sahraei S, Kalsoom Khan F, Desirò A, Bryson AE, Mondo SJ, et al. Evolutionary history of arbuscular mycorrhizal fungi and genomic signatures of obligate symbiosis. BMC Genomics. 2024;25(1):1–12. doi:10.1186/s12864-024-10391-2

59. Naranjo-Ortiz MA, Gabaldón T. Fungal evolution: major ecological adaptations and evolutionary transitions. Biol Rev. 2019;94(4):1443–76. doi:10.1111/brv.12510

60. Melie T, Pirro S, Miller AN, Smith SD, Schutz KS, Quandt CA. Comparative genomics and phylogenomic investigation of the class Geoglossomycetes provide insights into ecological specialization and the systematics of Pezizomycotina. Mycologia. 2023 Jul 4;115(4):499–512. doi:10.1080/00275514.2023.2186743

61. Baba T, Hagiuda R, Matsumae H, Hirose D. Does the genome of Sarcoleotia globosa encode a rich carbohydrate-active enzyme gene repertoire? Mycologia. 2025 Mar 4;117(2):255–60. doi:10.1080/00275514.2025.2452305

62. Green TGA, Smith DC. Lichen Physiology: Xiv. Differences Between Lichen Algae In Symbiosis And In Isolation. New Phytol. 1974;73(4):753–66. doi:10.1111/j.1469-8137.1974.tb01303.x

63. Patel TK, Williamson JD. Mannitol in Plants, Fungi, and Plant–Fungal Interactions. Trends Plant Sci. 2016;21(6):486–97. doi: 10.1016/j.tplants.2016.01.006

64. Richardson DHS. Photosynthesis and carbohydrate movement. In: The Lichens. Elsevier; 1973. p. 249–88. doi:10.1016/B978-0-12-044950-7.50013-1

65. Collins CR, Farrar JF. Structural resistances to mass transfer in the Lichen Xanthoria Parietina. New Phytologist. 1978;81(1):71–83. doi:10.1111/j.1469-8137.1978.tb01605.x

66. Komiya T, Shibata S. Polyols produced by the cultured phyco– and mycobionts of some Ramalina species. Phytochemistry. 1971;10(4):695–9. doi: 10.1016/S0031-9422(00)97135-8

67. Desai BM, Modi VV, Shah VK. Studies on polyol metabolism in Aspergillus niger. I. Nutritional requirements of a strain of Aspergillus niger cultivated on sorbitol as sole source of carbon. Arch Mikrobiol. 1969;67(1):6–11. doi: 10.1007/BF00413675.

68. Hallsworth JE, Magan N. Manipulation of intracellular glycerol and erythritol enhances germination of conidia at low water availability. Microbiology. 1995;141(5):1109–15. doi:10.1099/13500872-141-5-1109

69. Braun ML, Niederpruem DJ. Erythritol Metabolism in Wild-Type and Mutant Strains of Schizophyllum commune. J Bacteriol. 1969 Nov;100(2):625–34. doi:10.1128/jb.100.2.625-634.1969

70. Wright JR, Tourneau D Le. Utilization and Production of Carbohydrates by Pyrenochaeta terrestris. Physiol Plant. 1965;18(4):1044–53. doi:10.1111/j.1399-3054.1965.tb07003.x

71. Isenberg P, Niederpruem DJ. Control of erythritol dehydrogenase in Schizophyllum commune. Arch Mikrobiol. 1967;56(1):22–30. doi:10.1128/jb.100.2.625-634.1969

72. Nishimura K, Harada T, Arita Y, Watanabe H, Iwabuki H, Terada A, et al. Identification of enzyme responsible for erythritol utilization and reaction product in yeast Lipomyces starkeyi. J Biosci Bioeng. 2006;101(4):303–8. doi:10.1263/jbb.101.303

73. Carly F, Steels S, Telek S, Vandermies M, Nicaud JM, Fickers P. Identification and characterization of EYD1, encoding an erythritol dehydrogenase in Yarrowia lipolytica and its application to bioconvert erythritol into erythrulose. Bioresour Technol. 2018;247:963–9. doi:10.1016/j.biortech.2017.09.168

74. Niang PM, Arguelles-Arias A, Steels S, Denies O, Nicaud JM, Fickers P. In Yarrowia lipolytica erythritol catabolism ends with erythrose phosphate. Cell Biol Int. 2020;44(2):651–60. doi:10.1002/cbin.11265

75. Hill DJ, Smith DC. Lichen Physiology XII. The ‘Inhibition Technique’. New Phytol. 1972;71(1):15–30. doi:10.1111/j.1469-8137.1972.tb04806.x

76. Hoiczyk E, Hansel A. Cyanobacterial Cell Walls: News from an Unusual Prokaryotic Envelope. J Bacteriol. 2000;182(5):1191–9. doi: 10.1128/jb.182.5.1191-1199.2000

77. Pereira S, Zille A, Micheletti E, Moradas-Ferreira P, De Philippis R, Tamagnini P. Complexity of cyanobacterial exopolysaccharides: Composition, structures, inducing factors and putative genes involved in their biosynthesis and assembly. FEMS Microbiol Rev. 2009;33(5):917–41. doi:10.1111/j.1574-6976.2009.00183.x

78. Alhattab M, Kermanshahi-Pour A, Brooks MSL. Microalgae disruption techniques for product recovery: influence of cell wall composition. J Appl Phycol. 2019;31(1):61–88. doi:10.1007/s10811-018-1560-9

79. Baudelet PH, Ricochon G, Linder M, Muniglia L. A new insight into cell walls of Chlorophyta. Vol. 25, Algal Research. Elsevier B.V.; 2017. p. 333–71. doi:10.1016/j.algal.2017.04.008

80. Domozych DS, LoRicco JG. The extracellular matrix of green algae. Plant Physiol. 2024;194(1):15–32. doi:10.1093/plphys/kiad384

81. König J, Peveling E. Cell Walls of the Phycobionts Trebouxia and Pseudotrebouxia: Constituents and their Localization. Lichenol. 1984;16(2):129–44. doi:10.1017/S002428298400030X

82. Cordeiro LMC, Sassaki GL, Iacomini M. First report on polysaccharides of Asterochloris and their potential role in the lichen symbiosis. Int J Biol Macromol. 2007;41(2):193–7. doi:10.1016/j.ijbiomac.2007.02.006

83. González-Hourcade M, Braga MR, Del Campo EM, Ascaso C, Patinõ C, Casano LM. Ultrastructural and biochemical analyses reveal cell wall remodelling in lichen-forming microalgae submitted to cyclic desiccation-rehydration. Ann Bot. 2020;125(3):459–69. doi:10.1093/aob/mcz181

84. Takeda H, Nisizawa K, Miwa T. Histochemical and Chemical Studies on the Cell Wall of Prasiola japonica. Shokubutsugaku Zasshi. 1967;80(945):109–17. doi:10.15281/jplantres1887.80.109

85. Medwed C, Holzinger A, Hofer S, Hartmann A, Michalik D, Glaser K, et al. Ecophysiological, morphological, and biochemical traits of free-living Diplosphaera chodatii (Trebouxiophyceae) reveal adaptation to harsh environmental conditions. Protoplasma. 2021;258(6):1187–99. doi:10.1007/s00709-021-01620-6

86. West GS, Hood OE. The structure of the cell-wall and the apical growth in the genus Trentepohlia. New Phytol. 1911;10(7–8):241–9. doi:10.1111/j.1469-8137.1911.tb05594.x

87. Steinecke F. Pektosekappe und Schachtelbau bei Trentepohlia. Botanisches Archiv (Zeitschrift für die gesamte Botanik). 1929;24:525–530.

88. McCoy GA. Nutritional, morphological and physiological characteristics of Trentepohlia (I.U. 1227) in axenic culture on defined media [Thesis]. Corvallis: Oregon State University; 1978.

89. Hariharan GN, Krishnamurthy K V. Nature of the apical cap in Trentepohlia. Current Science. 1989;58(9)505

90. Good BH, Chapman RL. The Ultrastructure of Phycopeltis (Chroolepidaceae: Chlorophyta). I. Sporopollenin in the Cell Walls. Am J Bot. 1978 Jan;65(1):27. doi:10.2307/2442549

91. Brunner U, Honegger R. Chemical and ultrastructural studies on the distribution of sporopolleninlike biopolymers in six genera of lichen phycobionts. Can J Bot. 1985;63(12):2221–30. doi:10.1139/b85-315

92. Carbonero ER, Cordeiro LMC, Mellinger CG, Sassaki GL, Stocker-Wörgötter E, J. Gorin PA, et al. Galactomannans with novel structures from the lichen Roccella decipiens Darb. Carbohydr Res. 2005;340(10):1699–705. 1.

93. Nelsen MP, Plata ER, Andrew CJ, Lücking R, Lumbsch HT. Phylogenetic Diversity of Trentepohlialean Algae Associated with Lichen-Forming Fungi. J Phycol. 2011;47(2):282–90. doi:10.1111/j.1529-8817.2011.00962.x

94. Rindi F, Lam DW, López-Bautista JM. Phylogenetic relationships and species circumscription in Trentepohlia and Printzina (Trentepohliales, Chlorophyta). Mol Phylogenet Evol. 2009;52(2):329–39. doi:10.1016/j.ympev.2009.01.009

95. Zhu H, Hu Z, Liu G. Morphology and molecular phylogeny of Trentepohliales (Chlorophyta) from China. Eur J Phycol. 2017;52(3):330–41. doi:10.1080/09670262.2017.1309574

96. Zhu H, Hu Y, Liu F, Hu Z, Liu G. Characterization of the Chloroplast Genome of Trentepohlia odorata (Trentepohliales, Chlorophyta), and Discussion of its Taxonomy. Int J Mol Sci. 2019 Apr 10;20(7):1774. doi: 10.3390/ijms20071774.

97. Dědková K, Vančurová L, Muggia L, Steinová J. The plurality of photobionts within single lichen thalli. Symbiosis. 2025;95(1):35–63. doi:10.1007/s13199-025-01036-3

98. Kantnerová V, Škaloud P. The diverse world within: age-dependent photobiont diversity in the lichen Protoparmeliopsis muralis. FEMS Microbiol Ecol. 2025;101(11). doi:10.1093/femsec/fiaf096

99. Tagirdzhanova G, Saary P, Cameron ES, Allen CCG, Garber AI, Escandón DD, et al. Microbial occurrence and symbiont detection in a global sample of lichen metagenomes. PLoS Biol. 2024;22(11). doi: 10.1371/journal.pbio.3002862

100. Andrews S. FastQC: a quality control tool for high throughput sequence data. 2010. Available from: http://www.bioinformatics.babraham.ac.uk/projects/fastqc

101. Alneberg J, Bjarnason BS, De Bruijn I, Schirmer M, Quick J, Ijaz UZ, et al. Binning metagenomic contigs by coverage and composition. Nat Methods. 2014;11(11):1144–6. doi:10.1038/nmeth.3103

102. Nurk S, Meleshko D, Korobeynikov A, Pevzner PA. MetaSPAdes: A new versatile metagenomic assembler. Genome Res. 2017 May 1;27(5):824–34. doi:10.1101/gr.213959.116

103. Saary P, Mitchell AL, Finn RD. Estimating the quality of eukaryotic genomes recovered from metagenomic analysis with EukCC. Genome Biol. 2020;21(1):1–21. doi:10.1186/s13059-020-02155-4

104. Manni M, Berkeley MR, Seppey M, Zdobnov EM. BUSCO: Assessing Genomic Data Quality and Beyond. Curr Protoc. 2021;1(12):1–41. doi:10.1002/cpz1.323

105. Tagirdzhanova G, Saary P, Tingley JP, Díaz-Escandón D, Abbott DW, Finn RD, et al. Predicted Input of Uncultured Fungal Symbionts to a Lichen Symbiosis from Metagenome-Assembled Genomes. Genome Biol Evol. 2021;13(4):1–18. doi:10.1093/gbe/evab047

106. Palmer JM, Stajich J. Funannotate v1.8.1: Eukaryotic genome annotation. Zenodo; 2020. Available from: 10.5281/zenodo.1134477

107. Frith MC. A new repeat-masking method enables specific detection of homologous sequences. Nucleic Acids Res. 2011;39(4):e23. doi: 10.1093/nar/gkq1212.

108. Stanke M, Diekhans M, Baertsch R, Haussler D. Using native and syntenically mapped cDNA alignments to improve de novo gene finding. Bioinformatics. 2008;24(5):637–44. doi:10.1093/bioinformatics/btn013

109. Korf I. Gene finding in novel genomes. BMC Bioinformatics. 2004;5:59. doi:10.1186/1471-2105-5-59

110. Majoros WH, Pertea M, Salzberg SL. TigrScan and GlimmerHMM: two open-source ab initio eukaryotic gene-finders. Bioinformatics. 2004;20(16):2878–9. doi:10.1093/bioinformatics/bth315.

111. Haas BJ, Salzberg SL, Zhu W, Pertea M, Allen JE, Orvis J, et al. Automated eukaryotic gene structure annotation using EVidenceModeler and the Program to Assemble Spliced Alignments. Genome Biol. 2008;9(1):R7. doi:10.1186/gb-2008-9-1-r7

112. Lowe TM, Eddy SR. tRNAscan-SE: a program for improved detection of transfer RNA genes in genomic sequence. Nucleic Acids Res. 1997;25(5):955–64. doi:10.1093/nar/25.5.955

113. Jones P, Binns D, Chang HY, Fraser M, Li W, McAnulla C, et al. InterProScan 5: genome-scale protein function classification. Bioinformatics. 2014;30(9):1236–40. doi:10.1093/bioinformatics/btu031

114. Cantalapiedra CP, Hernández-Plaza A, Letunic I, Bork P, Huerta-Cepas J. eggNOG-mapper v2: functional annotation, orthology assignments, and domain prediction at the metagenomic scale. Mol Biol Evol. 2021;38(12):5825–9. doi:10.1093/molbev/msab293

115. Huerta-Cepas J, Szklarczyk D, Heller D, Hernández-Plaza A, Forslund SK, Cook H, et al. eggNOG 5.0: a hierarchical, functionally and phylogenetically annotated orthology resource based on 5090 organisms and 2502 viruses. Nucleic Acids Res. 2019;47(D1):D309–14. doi:10.1093/nar/gky1085

116. Mistry J, Chuguransky S, Williams L, Qureshi M, Salazar GA, Sonnhammer ELL, et al. Pfam: the protein families database in 2021. Nucleic Acids Res. 2021;49(D1):D412–9. doi:10.1093/nar/gkaa913

117. The UniProt Consortium. UniProt: the Universal Protein Knowledgebase in 2023. Nucleic Acids Res. 2023;51(D1):D523–31. doi:10.1093/nar/gkac1052

118. Yin Y., Mao X., Yang J., Chen X., Mao F., Xu Y. dbCAN: a web resource for automated carbohydrate-active enzyme annotation. Nucleic Acids Res. 2012; 40:W445–W451. doi:10.1093/nar/gks479

119. Rawlings ND, Barrett AJ, Thomas PD, Huang X, Bateman A, Finn RD. The MEROPS database of proteolytic enzymes, their substrates and inhibitors in 2017 and a comparison with peptidases in the PANTHER database. Nucleic Acids Res. 2018;46(D1):D624–32. doi:10.1093/nar/gkx1134.

120. Resl P, Hahn C. Phylociraptor: a unified computational framework for reproducible phylogenomic inference. bioRxiv. 2023. Preprint. doi:10.1101/2023.09.22.558970

121. Katoh K, Standley DM. MAFFT multiple sequence alignment software version 7: Improvements in performance and usability. Mol Biol Evol. 2013;30(4):772–780. doi:10.1093/molbev/mst010

122. Steenwyk JL, Buida TJ 3rd, Li Y, Shen XX, Rokas A. ClipKIT: a multiple sequence alignment trimming software for accurate phylogenomic inference. PLoS Biol. 2020;18(12):e3001007. doi:10.1371/journal.pbio.3001007

123. Minh BQ, Schmidt HA, Chernomor O, Schrempf D, Woodhams MD, von Haeseler A, Lanfear R. IQ-TREE 2: New models and efficient methods for phylogenetic inference in the genomic era. Mol Biol Evol. 2020;37(5):1530–1534. doi:10.1093/molbev/msaa015

124. Zhang C, Rabiee M, Sayyari E, Mirarab S. ASTRAL-III: polynomial time species tree reconstruction from partially resolved gene trees. BMC Bioinformatics. 2018;19:153. doi:10.1186/s12859-018-2129-y

125. Revell LJ. phytools 2.0: an updated R ecosystem for phylogenetic comparative methods (and other things). PeerJ. 2024;12:e16505. doi:10.7717/peerj.165

