## Supplementary figures and images for "Carbohydrate degradation machineries in lichen fungal symbionts reveal distinct symbiotic footprints across *Ascomycota*"

### Supplemental Figure

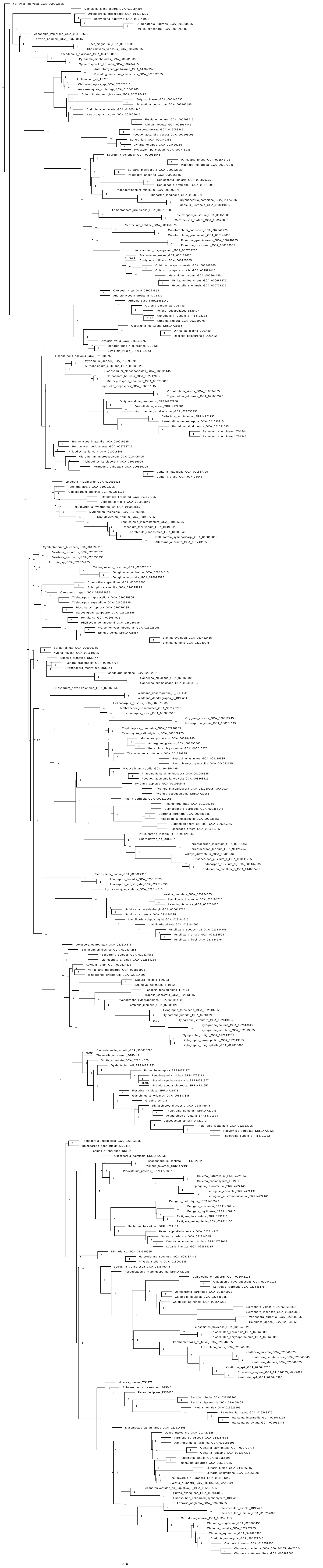

**Figure S1. Species tree from 309 fungal genomes across ascomycota based on 1387 gene trees.**
